# Framework for Martini-based Coarse-grained Model of Enzymes: Model Development and Experimental Validation

**DOI:** 10.1101/2024.09.22.614383

**Authors:** Mason Hooten, Sanjeeva Murthy, Nityananda Pal, Sagar D. Khare, Adam J. Gormley, Meenakshi Dutt

**Affiliations:** Biomedical Engineering, Rutgers, The State University of New Jersey, Piscataway, New Jersey 08854; Chemistry and Chemical Biology, Rutgers, The State University of New Jersey, Piscataway, New Jersey 08854; Chemical and Biochemical Engineering, Rutgers, The State University of New Jersey, Piscataway, New Jersey 08854

**Keywords:** Proteins, Enzymes, Coarse-grained models, Martini, Lipase, Dehalogenase, small-angle X-ray scattering

## Abstract

Recent experiments have shown that enzyme activity can preserved in harsh environments by complexing enzyme with polymer into a Protein-Polymer Hybrid (PPH). In a successful PPH, heteropolymer strands bind to the enzyme surface and restrain the folded protein without adversely affecting the binding and active sites. It is believed that hybridization is driven by non-covalent interactions at the enzyme surface including hydrophobicity and electrostatics. Molecular modeling of these interactions is not practical at the all-atom scale due to the long timescales and large particle counts needed to characterize binding. Protein structure at the scale of amino acid residues is parsimoniously represented by a coarse-grained model in which one particle represents several atoms, significantly reducing the cost of simulation. In this study we present two coarse-grained enzyme models – lipase and dehalogenase – prepared using a top-down modeling strategy. We simulate each enzyme in aqueous solution and calculate statistics of protein surface features and shape descriptors. The values from the coarse-grained data are compared with the same calculations performed on all-atom reference systems, revealing key similarities of surface chemistry at the two scales. Structural measures are calculated from the all-atom reference systems and compared with estimates from small-angle X-ray scattering (SAXS) experiments, with good agreement between the two. The described procedures of modeling and analysis comprise a framework for the development of coarse-grained models of protein surfaces with validation to experiment.

## 1. INTRODUCTION

Enzymes are proteins that act as biological catalysts. As they are naturally occurring, enzymes have several advantages over traditional industrial catalysts including high production efficiency, high substrate selectivity, and lower environmental impact [Chapman 2018]. They are used extensively in the areas of pharmaceuticals, food and beverage, and biofuel production [Chapman 2018]. Therefore, it is desirable to develop strategies for protein engineering which leverage enzyme function for industrial applications.

Enzyme catalytic activity is sensitive to the structural conformation of the protein itself. Techniques for stabilizing protein structure by covalent conjugation are likely to induce conformational changes which must also be controlled, increasing the complexity of protein design strategies [Keefe 2012; Panganiban 2018].

An alternative strategy is to induce stability via noncovalent adsorption of random heteropolymers to form protein polymer hybrids (PPHs) [Chapman 2019; Lancaster 2018; Pelegri-O’Day 2014; Ko 2018; Kosuri 2022]. The PPH approach has demonstrated the capacity to retain high catalytic activity in stress environments [Panganiban 2018]. However, chemical theories to guide polymer composition for targeted PPH formation are nascent. Robotic platforms have been demonstrated to characterize candidate combinations of enzymes and heteropolymers [Tamasi 2022], potentially reducing the cost of developing viable PPH chemistries given a constrained polymer design space. However, this approach is still limited in the size of a candidate polymer library that may be thoroughly investigated, and therefore there is little reason to believe that experimental results will generalize between proteins.

Computational models provide a means for exploration of a larger polymer design space with lower cost and higher precision than what is possible on the bench. Particle-based models coupled with Molecular Dynamics (MD) simulations enable investigation of the chemically-specific dynamics which govern the coassembly of active enzymes with adjunct polymers via surface adsorption. Analysis of the emergent physics from these MD simulations may significantly reduce the size of viable candidate libraries for experimental validation, or may allow further theoretical generalization of the PPH design task as a function of protein and polymer properties.

The general dynamics by which biomacromolecules dock with synthetic monomers are inaccessible to all-atom MD simulations due to the large spatiotemporal scales of multicomponent assembly. Coarse-grained models overcome this limitation by judiciously grouping atoms to be represented instead by pseudo atoms, or coarse-grained beads. The coarse-grained representation therefore increases computational efficiency at the cost of increased uncertainty in atomic positions.

Another key assumption in coarse-grained models of proteins is that the chosen scale of the coarse-grained beads will provide resolution of all relevant structural changes in a dynamical simulation. This assumption is encoded in protein models by choosing bead definitions which group atoms at a smaller scale than the proteinogenic amino acids, therefore preserving some intra-residue conformational flexibility.

The validity of these assumptions cannot be tested via experimental methods. Hence, the models are calibrated via structural measurements which may be compared with corresponding experimental results. A multiscale approach is adopted to validate the coarse-grained model. In the first step, structural calculations from a reference all-atom model are compared with data from X-ray diffraction (XRD) experiment. Small-angle X-ray scattering (SAXS) may be used to measure the structure function, S(q), and estimate the pair-distance distribution function (PDF) of an enzyme in aqueous solution, while the PDF may be calculated directly from MD trajectories. This step is followed by comparison of structural and surface features of the target coarse-grained model with the all-atom reference.

Many biomolecular phenomena are assumed to be largely mediated by molecular interfaces, including the surfaces of individual molecules in solution. It is therefore assumed that surface composition would be a powerful predictor of PPH formation. While direct experimental determination of protein surfaces in solution is unfeasible, the validated all-atom reference model provides extensive data for comparison. For the validation of the secondary model – coarse-grained versus all-atom – a set of structure and surface descriptors are selected to summarize the ensemble behavior of the enzyme models.

Particle-based models typically comprise a choice of structural representation and a force field. Several popular strategies for the development of coarse-grained MD models of biomolecules have been developed, with a correspondent variety in their advantages and development costs [Banerjee 2024]. In bottom-up approaches, pseudo atoms represent specific constituent atoms and potential energy functions are derived using simulated all-atom data. Such derivation often is either the parameterization of specific functional forms, such as a Lennard-Jones potential, or else a direct estimate of the potential energy surface via tabulated values. This approach to representation allows for arbitrarily high chemical specificity of the resulting model, with the coarse-grained force fields able to recapitulate the objective features such as the forces, entropy, or structures of the reference set. Parameterization utilizes techniques such as iterative Boltzmann inversion, force matching, relative entropy and inverse Monte Carlo [Reith 2003; Rühle 2011; Lyubartsev 1995; Shell 2008]. Frameworks built using these techniques have been used to resolve conformational behavior and self-assembly of biomolecules and biomimetics including peptides [Izvekov 2006, Carmichael 2012; Villa 2009; Hooten 2023; Banerjee 2022; Banerjee 2023; Hooten 2025 [in preparation] ].

Bottom-up approaches typically require extensive amounts of chemically specific reference data, which puts a practical limit on model transferability. More recently, bottom-up coarse-grained models have been parameterized with machine-learned (ML) potentials [Durumeric 2023; Ruff 2015; Sahrmann 2023]. Depending on the architecture, some of these models are capable of implicitly including numerous many-body interactions, transcending one of the traditional engineering constraints in which many-body behavior is approximated using only two-body potentials. It remains an open question whether ML potentials can produce force fields which are transferable over large molecular design spaces – like general protein folding – if appropriate reference data is available [Durumeric 2023]. However, the data-intensive nature of the modern ML approach may not be suitable where a very large or even an open-ended design space is considered.

Alternatively, models may be designed to target larger scale phenomena, and calibrated using data such as liquid-liquid partitioning or system free energy obtained from experiment or theory [Banerjee 2024; Mushnoori 2018; Mushnoori 2023; Aydin 2014; Aydin 2016; Chong 2016; Muthukumar 2015]. These models are based on methods classified as top-down approaches. Top-down models typically employ coarse-grained bead definitions and resolutions intended to express the general character of molecular structure while de-emphasizing specific chemical features of the atomistic representation. For example, the Martini coarse-grained framework has been applied to investigate self-assembly of biomolecular nanostructures including lipid bilayers [Marrink 2007; Marrink 2019] and large supramolecular assemblies of proteins and lipids [Reddy 2016; Wang 2022], as well as to probe the binding and diffusion characteristics of small molecules in contact with membrane bilayers [Li 2018].

For this study, coarse-grained models of proteins are developed using Martini 3 [Souza 2021]. Martini is a popular framework for development of coarse-grained particle-based models of biomolecules and polymers. It provides a standard set of coarse-grained beads defining size, polarity, and ionic character, as well as a library of standard structures for the proteinogenic amino acids. Thus, Martini provides a good foundation for the bootstrapping of a variety of protein models.

In this study, two enzyme systems, lipase and dehalogenase, are investigated using a combination of MD simulations at all-atom and coarse-grained resolutions, along with SAXS experiments on commercially available proteins for comparison. The all-atom MD simulations of proteins are performed using the CHARMM force field, chosen for its ability to recapitulate folded protein structure [Huang 2017]. Ensembles of all-atom MD simulations are compared to SAXS measurements via the unnormalized PDF and radius of gyration (*R_g_*). Distributions of the gyration tensor parameters are calculated for simulated structures at all-atom and coarse-grained resolutions and the results are compared. Surface compositions are also calculated using the SURFMAP molecular cartography software at all-atom and coarse-grained resolutions and the results are compared.

The salient features of the gross structure observed in experiment and expressed by *R_g_* and PDF are captured in the all-atom model. While this indicates that the comparison of all-atom to SAXS is reasonable, the experimental preprocessing is found to be a critical step to control for the presence of excipients or other off-target species which may be present in the sample. Calculated shape and surface features fluctuate over similar ranges at both all-atom and coarse-grained resolution, indicating qualitative agreement between the models. The surface composition of the all-atom and coarse-grained models is found to be similar in the distributions of characteristics of surface features, owing to the contributions of individual residues.

Overall, the study demonstrates the feasibility of a multiscale validation design strategy for the preparation of coarse-grained models. A framework encompassing a process and a set of benchmarks is presented for the development and validation of coarse-grained enzyme models using this multiscale approach. These findings open the door to new protein engineering design strategies based on enzyme surface features.

## 2. METHODS

### 2.1 All-Atom Molecular Dynamics Representation of Enzymes and Simulations

All-atom MD simulations were performed using Gromacs 2021 [Abraham 2015; Berendsen 1995; Lindahl 2001; Van Der Spoel 2005] to provide appropriate reference data. A single protein molecule is simulated in a cubic box, with three-dimensional periodic boundary conditions using the CHARMM36m force field [Huang 2016]. CHARMM was chosen because it is parameterized to be suitable to study the dynamics of protein conformation, with attention given to globular proteins as well as intrinsically disordered proteins [Best 2012; Huang 2016]. The edge length of the simulation box is equal to the diameter of the molecule plus 3 nm, adhering to the minimum image convention to prevent self-interaction of atoms [Allen 2017a].

An initial lipase reference structure was derived from PDB [Berman 2000] structure 1OIL [Kim 1997]. Gromacs is used to transform the PDB structures to the native Gromacs structure format. Scripts with details of all Gromacs commands used are available on the public GitHub repository supporting this study [Hooten 2024]. Systems are solvated with 30,481 water molecules using the TIP3P explicit water model [Jorgensen 1983]. Charge neutralization is achieved by the inclusion of a single calcium cation which was present in the PDB structure along with 2 sodium cations. Details of system composition may be found in Table S1 in the Supporting Information file.

Dehalogenase systems were created using an initial structure derived from PDB structure 3RK4 [Lahoda 2014]. Systems are solvated with 23,166 water molecules. Charge is neutralized by inclusion of a single chloride anion which was present in the PDB structure along with 18 sodium cations.

All-atom MD simulations were performed to explore the conformation of the biomolecule in aqueous solution. Detailed simulation parameters may be found in Table S2 in the Supporting Information file, as well as in the GitHub repository [Hooten 2024]. Solvated enzyme systems are first subject to steepest descent energy minimization until the maximum force falls below 1000 kJ mol-1 nm-1. Energy minimization is followed by an NVT equilibration step in which the atoms constituting the protein are restrained to allow relaxation of the solvent. Equilibration temperature is set to 300 K using the V-rescale thermostat with a stochastic term [Bussi 2007]. Electrostatic forces are calculated using the Particle Mesh Ewald (PME) method [Darden 1993] with a 1.0 nm cutoff. The system is simulated using the leapfrog integrator for a simulation spanning 1 ns, using a time step of 2 fs.

In production simulations, the temperature is fixed at 300 K using the Nosé-Hoover thermostat [Nosé 1984; Hoover 1985], and the pressure is fixed at 1 bar using the Parrinello-Rahman barostat [Parrinello 1981], both with a dispersion correction applied. Electrostatic forces are calculated using PME [Darden 1993] with a 1.0 nm cutoff. Dispersion forces are calculated using a 1.0 nm cutoff. Periodic boundaries are applied in the three directions. The box length is chosen such that the distance between any atom and the box boundary is greater than the cutoffs of the potential. The leapfrog integrator is used with a time step of 2 fs. Ten independent trajectories per enzyme sequence were simulated for durations spanning 500 ns, yielding 5 microseconds of cumulative simulated time per sequence. Production simulations were performed using the Pittsburgh Supercomputing Center Bridges-2 cluster [Brown 2021; Towns 2014].

Trajectories of all atoms were sampled at 2 ps intervals and subsequently resampled for analysis. For calculations of interatomic pair distances, trajectories including only the heavy (i.e., non-hydrogen) atoms were used. Each independent trajectory was sampled over its entire 500 ns duration at intervals of 50 ps, yielding 10,001 samples per trajectory and 100,010 samples per enzyme sequence. Root mean square fluctuation (RMSF), gyration tensor shape parameters, and surface properties were calculated from all-atom trajectories including only the heavy (non-hydrogen) atoms of the enzyme. RMSF and the shape parameters are sampled from 400 to 500 ns at intervals of 10 ps (which translates to 10,001 samples per independent trajectory, 100,010 samples per ensemble). Surface properties are sampled over the same span but at intervals of 10 ns (which translates to 11 samples per independent trajectory, 110 samples per ensemble).

### 2.2 Coarse-grained Molecular Dynamics Representation of Enzymes and Simulations

The configurational complexity of proteins, especially in high-resolution representations, means that deriving a force field which maintains a protein in a globular conformation remains an open problem. In coarse-grained representation of proteins, methods have been devised to systematically maintain tertiary structure in proteins by damping or constraining the pairwise interaction of atoms which are not covalently bonded [Banerjee 2024; Poma 2017; Tirion 1996; Bahar 1997].

One common strategy is to extend the typical bead-spring model with an auxiliary set of nonbonded pair interactions between nearest neighboring atoms within the protein which are parameterized to bias for native conformation, the so-called Go-like approach. Go-like models have the advantage of allowing unconstrained dynamics, although the use of many additional pair potentials may substantially increase the cost of computation [Poma 2017].

A popular alternative is to add unbreakable harmonic potentials to atoms whose pairwise distance falls within some target range, so that the covalently bonded structure of the protein is supplemented with further constraints on the relative motion of distant pairs of atoms, effectively maintaining its tertiary structure. A model which includes such a set of long-distance harmonic constraints is referred to as an elastic network model. Such elastic network models have been used in computational investigations of proteins which probe the dynamics while maintaining a critical assumption that the protein is in its native structure.

Some early applications of elastic network models of proteins apply force fields consisting only of an elastic network, while excluding both the more conventional Lennard-Jones type nonbonded interactions [Tirion 1996; Bahar 1997] and even excluding the explicit parameterization of the covalently bonded chain [Tirion 1996]. For example, coarse-grained models with elastic networks have been used to elucidate the dynamics entailing the binding of small-molecules in large proteins or protein complexes [Souza 2020; Gutiérrez-Fernández 2020; Coupland 2021; Pezeshkian 2023]. Other investigators have used coarse-grained models with elastic networks to examine membrane-binding processes, enabling the identification of particular regions of a protein which are involved in the initial association with the membrane [Synek 2021]. Further details regarding applications and analysis of elastic network models are reviewed in reference [Togashi 2018].

Although the strength of the harmonic bond may be tuned to modulate the flexibility of the system, an elastic network model fundamentally limits system dynamics and therefore may not be suitable for applications where conformational changes are of interest. A coarse-graining framework with an elastic network, which employs all-atom configurations as a reference, will use a highly constrained network. The configurations sampled using a coarse-grained model with an elastic network should hypothetically mimic corresponding configurations sampled using all-atom models, thereby providing an optimal benchmark for comparison between the two resolutions. Hence, this study adopts a coarse-graining framework based upon the Martini and elastic network models.

In this study, the tertiary structure of the protein is maintained by overlaying an elastic network of pseudobonds upon the coarse-grained representation of the molecule. The network adds a harmonic potential between all coarse-grained bead pairs at distance between 0 and 0.9 nm with a bond strength of 500 kJ mol-1 nm-2. The coarse-grained Martini 3 force field (version 3.0.0 parameter set [Souza 2021]) was used to represent the amino acids and capture their mutual interactions.

Explicit solvent, coarse-grained MD simulations of a single protein molecule solvated in water were performed. For each independent trajectory, an all-atom configuration is transformed to a coarse-grained configuration using the Martinize2 program [Kroon 2022] to provide an initial structure. The coarse-grained MD simulations are initialized by using a box length that corresponds to the one from the coarse-grain-mapped all-atom reference configuration. Figure 1 shows schematic images of lipase and dehalogenase, contrasting their all-atom and coarse-grained representations and visualizing the modeled elastic networks.

**Figure 1.**
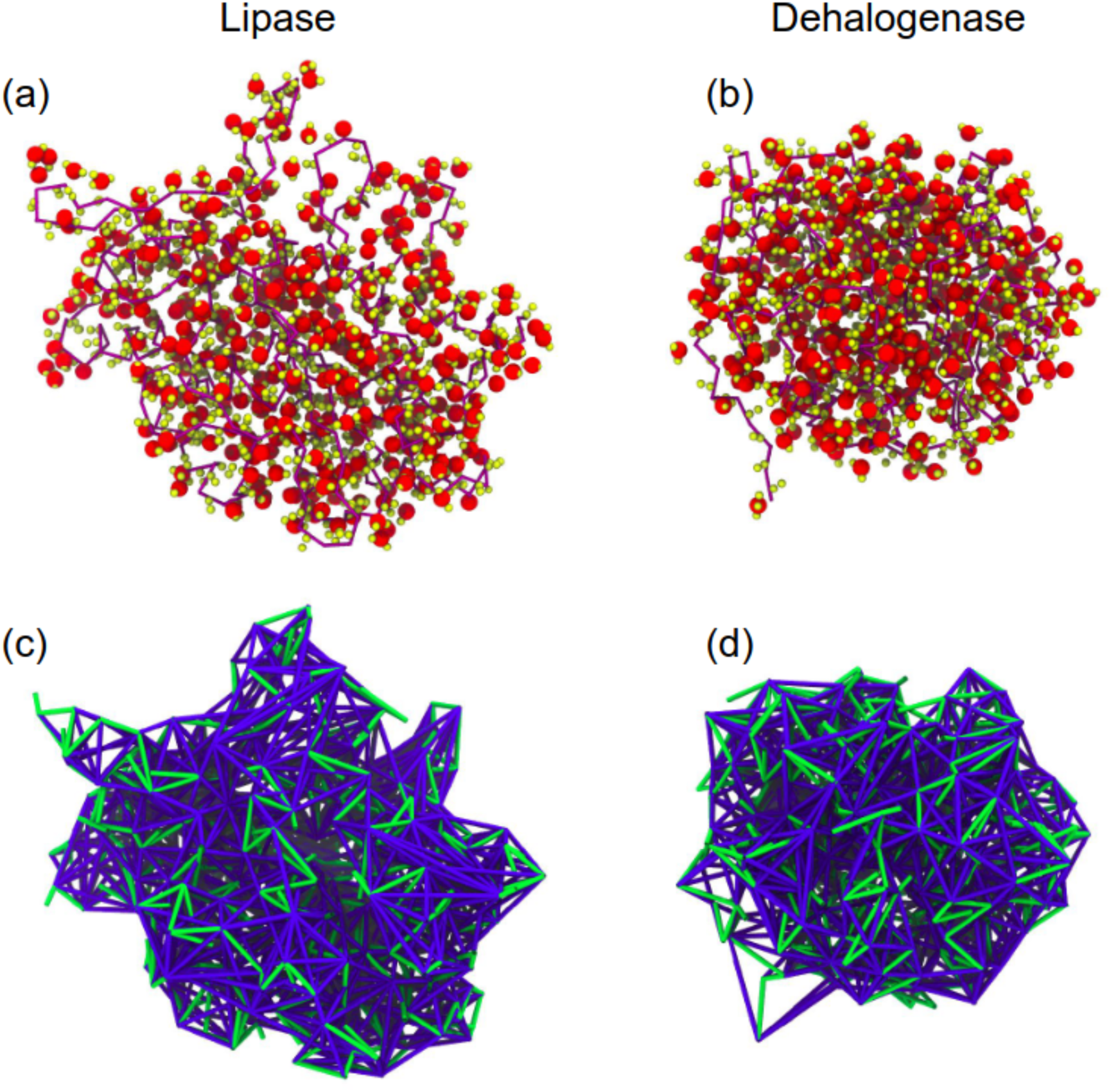
Molecular visualizations comparing coarse-grained and all-atom representations. Upper panels (a) and (b) show contrast between individual particle counts at each resolution, in models of lipase and dehalogenase, respectively. The tortuous purple line traces the alpha carbons of the all-atom backbone, with close analogy to the backbone beads in the coarse-grained representation. Yellow spheres represent the other non-hydrogen atoms of the all-atom representations, where hydrogen atoms constitute roughly half of the total number of atoms in each protein. Red spheres represent the coarse-grained beads, excluding the backbone beads. Lower panels (c) and (d) show the bond structure of the coarse-grained representations. Green lines represent the bonds between coarse-grained beads, blue lines represent the auxiliary pseudobonds of the elastic network.

Prior to solvation, the coarse-grained representations of the enzyme systems in vacuum are subject to steepest descent energy minimization until the maximum force falls below 1000 kJ mol-1 nm-1. The systems are then solvated and subject to a second round of energy minimization. Each system is then subject to a three-part equilibration routine. First, the enzyme is restrained and 2,000 steps of NVT simulation are performed using a time step of 5 fs. Next, the restraints are removed and an additional 25,000 steps of NVT are performed using a time step of 2 fs. Then a pressure correction is applied and the system is simulated for 10,000 steps of NPT at a time step of 10 fs. The reasoning underlying the extended routine is to first allow the solvent to relax, then allow the enzyme to adjust to local solvent changes before slowly accelerating dynamics. All equilibration simulations use the leapfrog integrator. The temperature is maintained at 300 K via the V-rescale thermostat with a stochastic term [Bussi 2007]. Calculations of electrostatic interactions use PME with a cutoff of 1.0 nm. During NPT equilibration, pressure is maintained at 1 bar using the Berendsen barostat [Berendsen 1984].

In production simulations, the temperature is fixed at 300 K using the Nosé-Hoover thermostat, and the pressure is fixed at 1 bar using the Parrinello-Rahman barostat, both with a dispersion correction applied. The electrostatic interactions are calculated using PME with a 1.1 nm cutoff. The dispersion forces are calculated with a 1.1 nm cutoff. The cubic simulation box has periodic boundaries. The leapfrog integrator is used with a simulation step of 20 fs. Ten independent coarse-grained trajectories per enzyme sequence were simulated for 25 million iterations, yielding 250 million total steps or 5 microseconds of cumulative simulated time per sequence.

Trajectories of all pseudoatoms were sampled at 2 ps intervals and subsequently resampled for analysis. RMSF was calculated from coarse-grained trajectories including only the backbone beads of the protein sampled from 100 to 500 ns at intervals of 40 ps (which translates to 10,001 samples per independent trajectory, 100,010 samples per ensemble). Gyration tensor shape parameters and surface properties were calculated from coarse-grained trajectories including all coarse-grained beads in the protein. Shape parameters are sampled from 100 to 500 ns nominal simulation time at intervals of 40 ps (which translates to 10,001 samples per independent trajectory, 100,010 samples per ensemble). Surface properties were sampled over the same time span but at intervals of 40 ns (which translates to 11 samples per seed, 110 samples per ensemble).

### 2.3 Characterization of Structure of Enzymes

#### Pair-distance Distribution Function

The PDF, P(r), is a measure of the distances between heavy atoms within an enzyme or protein. This quantity may be calculated directly from simulation data [Allen 2017; Allen 2017b]. The PDF may be estimated at different scales by means of XRD techniques, with SAXS used for long interatomic distances (>10 Å) and wide-angle x-ray scattering (WAXS) for short distances (<10 Å). Here we will confine ourselves to PDFs obtained from SAXS data.

### Shape Characterization, Gyration Tensor Parameters

The structure of a protein is the fundamental determinant of its function. In this study, shape descriptors derived from the gyration tensor *S*, given by equation (1), are used to characterize the structure of the simulated enzymes. The gyration tensor captures the mass distribution of a set of particles, and its associated shape descriptors have been demonstrated as a means of distinguishing the sphericity and compactness in a variety of coarse-grained representations of polymers and proteins [Arkin and Janke 2013; Wang 2012].

The three shape descriptors that are computed are invariants determined from the gyration tensor, as given by equations (2) to (4). The *R_g_* is a measure of the rotational inertia, which is correlated with chain length in globular proteins [Dima 2004]. The relative shape anisotropy (*κ^2^*) measures the shape symmetry, ranging from 0 for highly symmetric objects to 1/4 for a square planar array of particles to 1 for linear or rod-like arrangement of particles. The asphericity (b) measures the deviation from spherical symmetry, which equals zero for spherically symmetric objects (including spheres and tetrahedrons), or greater than zero otherwise [Theodorou 1985]. Implementations of these calculations may be found in the public GitHub repository [Hooten 2024].

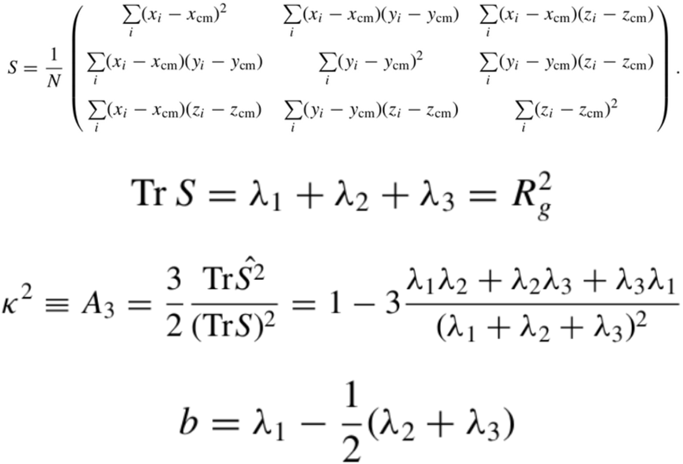

Equations 1 thru 4. Definition of the gyration tensor S, and definitions of its invariant shape descriptors *R_g_*, *κ^2^*, and *b*.

2.4 **Methods - X-ray Diffraction Measurements**

#### 2.4.1 Materials

Burkholderia cepacia (formerly Pseudomonas cepacia) lipase, an extensively studied lipase, was used in these experiments. Crude lipase (Amano, Sigma Aldrich; MW 33 kDa) that was purchased was found to contain < 1 wt. % of lipase, the rest being a stabilizer (starch; Cassava dextrin 100-250 kDa, private communication, Sigma Aldrich; confirmed by Fourier Transform Infrared Spectroscopy, FTIR). This lipase was purified using the methods described in the literature [Bornscheuer 1994; Hedrich 1991], and was concentrated by using ultrafiltration spin columns [Luić 2001]. The concentrations were measured using a Nanodrop instrument. Purity of the enzyme was confirmed by sodium dodecyl sulfate–polyacrylamide gel electrophoresis (SDS-PAGE) analysis. Activity was measured using 4-Nitrophenyl Butyrate (PNB) substrates (Sigma-Aldrich).

Haloalkane dehalogenase (DhaA31) enzyme was prepared following Ref(ChemBioChem 2017, 18, 2000–2006) with several modifications. DhaA31 was overexpressed in competent cells of the *E. coli* BL21 strain, which were transformed with a pET29b+ vector containing the DhaA31 gene (Twist Biosciences) with a C-terminal 6xHis-tag. The transformed cell line was grown in 500 ml of ZY auto-induction media with a 5 mL preculture at 37°C for 3 hours until the optical density reached 1.8–2. The temperature was then lowered to 18°C after 3h, and further incubated at 18C for 21 hours. Cells were collected via centrifugation at 5000 rpm for 40 minutes and resuspended in 35 mL of wash buffer (50 mM Na2HPO4, 500 mM NaCl, 30 mM imidazole, pH 7.4). 40 μL of 1 M PMSF, lysozyme, and DNase (final concentrations of ∼2 mg/mL and ∼0.2 mg/mL, respectively) were added to the resuspension. The resuspended cells were lysed via sonication on ice, and the lysate was centrifuged at 70,000 rpm for 40 minutes. The DhaA31 enzyme containing 6xHis-tag was purified using Ni-NTA columns (Qiagen) and eluted with 8 mL of elution buffer (50 mM Na2HPO4, 500 mM NaCl, 500 mM imidazole, pH 7.4). The eluate was dialysed against 0.1 M phosphate buffer (pH 7.4) to remove the excess imidazole. Enzyme purity was analyzed by SDS-PAGE (see Supporting Information, Figure S4 for SDS-PAGE data) and characterized by mass spectrometry (see Supporting Information, Figure S5 for mass spectrometry data). The enzyme concentration was measured using a NanoDrop (DS-11+, DeNovix). The yield of the enzyme was 157 mg/L. Enzyme activity was measured following Ref (ChemBioChem 2017, 18, 2000–2006) with 1-chlorobutane as the substrate.

#### 2.4.2 Small-Angle X-ray Scattering

SAXS experiments were conducted at beamline 16-ID for Life Science X-ray Scattering (LiX; National Synchrotron Light Source II, Brookhaven National Laboratory, Upton, NY).

SAXS data were collected with 15.14 keV X-rays and three Pilatus 1M detectors collected over a *q* range of 0.005-3.13 Å^-1^. Background subtraction was done using the buffer scattering obtained after every three samples, and by scaling to the water peak at q = 2 Å^-1^. Data at *q* values of 0.005-0.25 Å^-1^ was used for further analysis. The data were analyzed in BioXTAS RAW 2.1 with ATSAS 3.0.4 to determine the *R_g_* by Guinier analysis. Pair-distance distribution function, P(r), was obtained by indirect Fourier transform with GNOM [Hopkins 2017; Hopkins 2020; Hopkins 2018; Manalastas-Cantos 2021; Franke 2017]. Bead model reconstructions using a dummy atom model were created with the collected SAXS data as input. These models were generated using the DAMMIN program in the ATSAS 3.0.4 software package assuming a single-phase object [Svergun 1999]. a file that was fed into DAMMIN for final refinement in slow mode using default parameters to improve model fitting [Volkov 2003].

Lipase measurements were carried out in potassium phosphate pH 7.4 at concentrations from 0.8 to 2.4 mg/mL. The *P(r)* data used here was obtained from a 1.2 mg/mL solution. Dehalogenase measurements were carried out in potassium phosphate buffer (pH ∼ 7.5) at 8.2, 3.9 and 0.9 mg/ml concentration. *P(r)* data used here was generated using the data from the 8.2 mg/mL solution.

## 3. Results and Discussion

### 3.1 Comparison of Results from All-Atom Trajectories and X-ray Diffraction Measurements

The *R_g_* from various sources of both the proteins are listed in Table 1. The measured *R_g_* of lipase is 22.4 Å. The only structural data available in the literature for psuedomonas cepacian lipase gives the dimensions of the globular dimensions derived from the crystal structure as 30 x 40 x 50 Å (Kyeong Kyu Kim et al. Structure 1997, Vol 5 173-185), from which the *R_g_* can be calculated to be 19.1 Å. The large SAXS can be attributed to the hydration shell around the proteins.

**Table 1.** Radii of gyration in Å of simulated and experimental systems. Averages reported for simulated values. * Lipase XRD value is calculated from PDB structure 1OIL.

|  | AA MD | SAXS | Literature |
| --- | --- | --- | --- |
| Lipase | 19.8 | 22.4 | 19.1* |
| Dehalogenase | 18.2 | 23.7 | 18.3 |

**Table 2.** Gyration tensor parameter calculations for models of Lipase and Dehalogenase at all-atom and coarse-grained resolution. For all-atom representations, only heavy atoms are included in the calculations. All beads are considered in the coarse-grained representations.

| Simulated System | Rg |  |  | K<sup>2</sup> |  |  | b |  |  |
| --- | --- | --- | --- | --- | --- | --- | --- | --- | --- |
|  | Mean | 25 percentile | 75 percentile | Mean | 25 percentile | 75 percentile | Mean | 25 percentile | 75 percentile |
| Lipase, all-atom | 1.977 | 1.955 | 1.989 | 0.078 | 0.069 | 0.086 | 1.061 | 0.986 | 1.133 |
| Lipase, coarse-grained | 1.898 | 1.890 | 1.907 | 0.032 | 0.028 | 0.035 | 0.624 | 0.588 | 0.661 |
| Dehalogenase, all-atom | 1.817 | 1.809 | 1.824 | 0.025 | 0.022 | 0.027 | 0.469 | 0.437 | 0.496 |
| Dehalogenase, coarse-grained | 1.777 | 1.77 | 1.784 | 0.024 | 0.022 | 0.026 | 0.414 | 0.387 | 0.441 |

The measured *R_g_* of DhaA31 is 23.7 Å. There is no structural data available in the literature with which to compare with the results reported in this study. One report with SAXS data from DhaA115 finds a much smaller *R_g_* than the corresponding result reported in this study (18.3 Å), but shows a scattering curve with features similar to ours [Markova 2020].

Figure 3 shows the pair-distance distribution function (PDF), *P(r),* derived from SAXS data for both lipase and DhaA31. Overlaid on these are curves calculated from all-atom data for both lipase and dehalogenase for comparison. It can be seen that the all-atom calculated distributions and the SAXS distributions typically achieve their peak values at a distance of around 24 Å. The majority of the distribution mass is typically concentrated in a segment with a nearly Gaussian shape, which is typical of globular macromolecules and indicates the presence of a well-established monomeric form of each enzyme system. The all-atom distribution also includes several spikes at short distances, from 1 to 5 Å, indicating the presence of regular structural units of consistent length, e.g., the length of the repeating peptide bond.

The maximum distances (*D_max_*) found in the calculated monomeric structures are 60 Å for lipase and 50 Å for DhaA31. *D_max_* could not be accurately determined from experimental data because of the long tails. The tails at larger distances indicate the presence of some extended conformation or some oligomeric species. The secondary peak at 60 Å, weak in lipase and strong in Dehalogenase is most likely due to the presence of dimers. Such dimers have been postulated both for lipases [Rios 2018] and for dehalogenases [Ridder 1999; de Jong 2003]. Despite the presence of the dimeric species, it can be seen there is a very good agreement between the calculated and measured *D_max_* values with lipase in which the dimer’s contribution is weak; it is difficult to assess this with DhaA31 because of the large contribution to the intensity from the dimers. Calculations using simple models with dimers constructed as spheres in contact indeed show that the extra shoulder and larger *D_max_* seen in the experimental data can be attributed to some dimers present in the solution (SI, Figure S7). While the crystallographic structure of DhaA31 (PDB:3RK4) reports the protein as a monomer, dimeric species have been reported in the literature for other haloalkane dehalogenase variants: DhaA115 Type 1 (PDB:6TY7), DhaA115 Type 2 (PDB: 6XT8), and DhaA177 (PDB:6XTC) [Markova 2021]. The existence of domain-swapped dehalogenase dimers in solution has also been confirmed experimentally by SAXS and cross-linking coupled mass spectrometry (XL-MS) data [Markova 2021]. Furthermore, a dimeric form was also observed in the crystal structure of a closely related DhaA variant (PDB:4KAF) [RCSB Protein Data Bank 2024]. These observations suggest that a significant amount of dimer may exist in the solution and contribute to the observed peak in the PDF.

The almost complete overlap of the PDFs of the two enzymes reflects the similarity between the dominant monomeric species of the two enzymes. The molecular weights of the two enzymes are very similar (33128 Da for lipase and 34189 Da for DhaA31), and both enzymes have similar architecture: they belong to the α/β-hydrolase fold family, and have two domains - a Rossman-fold domain with a pattern of alternating alpha helices and beta strands forming a parallel β-sheet, and a majority helical lid domain, as shown in Figure 2. The active site is situated at the domain-domain interface with a similar spatial positioning of the catalytic triad residues [Nardini 1999]. This conserved topology means that on average residue pair distances are likely to be similarly distributed resulting in similar PDFs.

**Figure 2.**
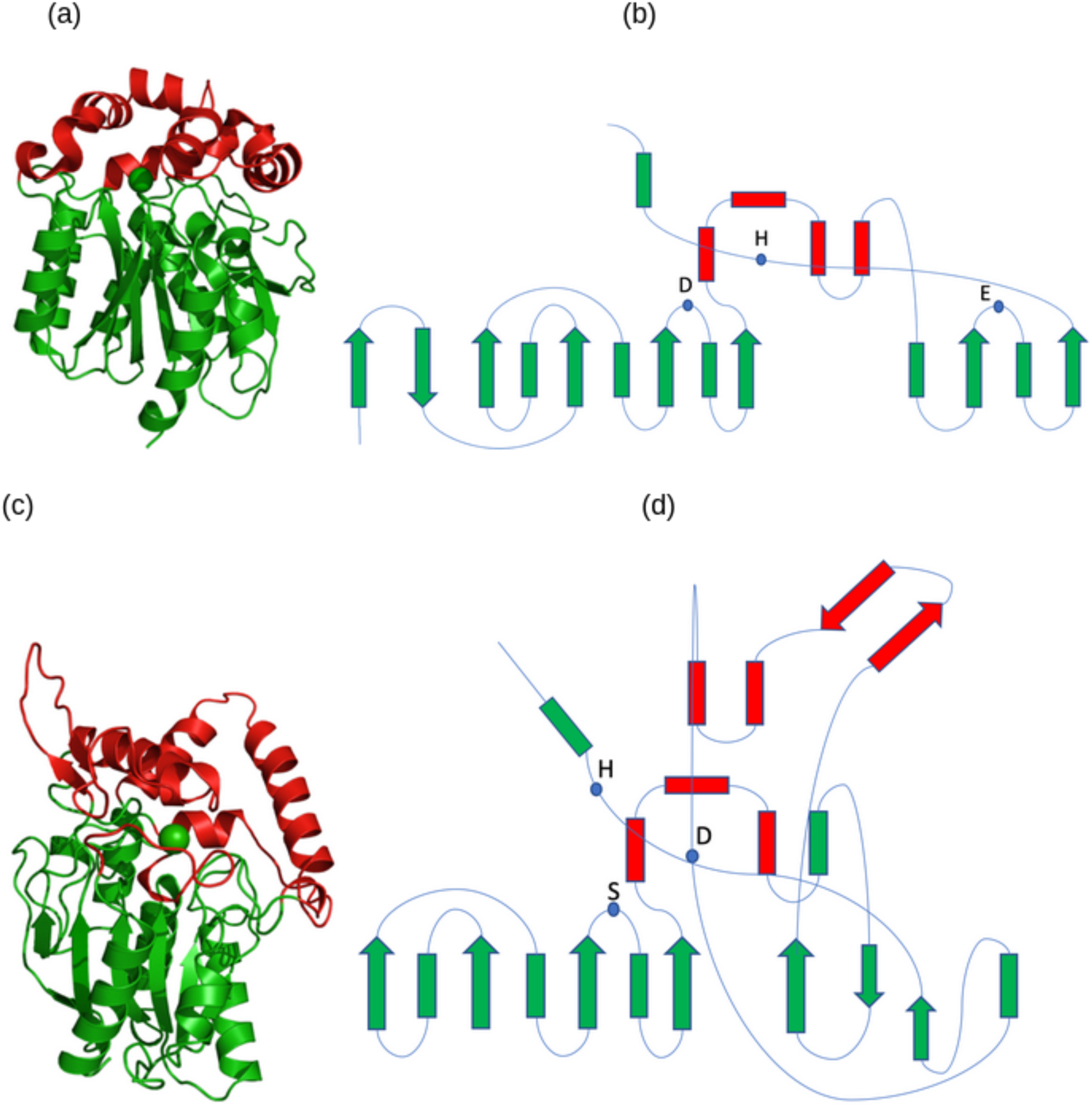
(a) 3D structure, and (b) topology diagram of the haloalkane dehalogenase DhaA31 (PDB:3RK4). The lid domain is shown in red and the Rossman fold domain is shown in green. α Helices and β strands belonging to the ‘Rossman’ fold domain (lid domain) are represented by green (red) rectangles and green (red) arrows, respectively. The location and identity of the catalytic triad is indicated by blue dots. Both the (c) 3D structure and (d) topology diagram of the lipase (PDB:1OIL) are highly similar to the dehalogenase.

**Figure 3.**
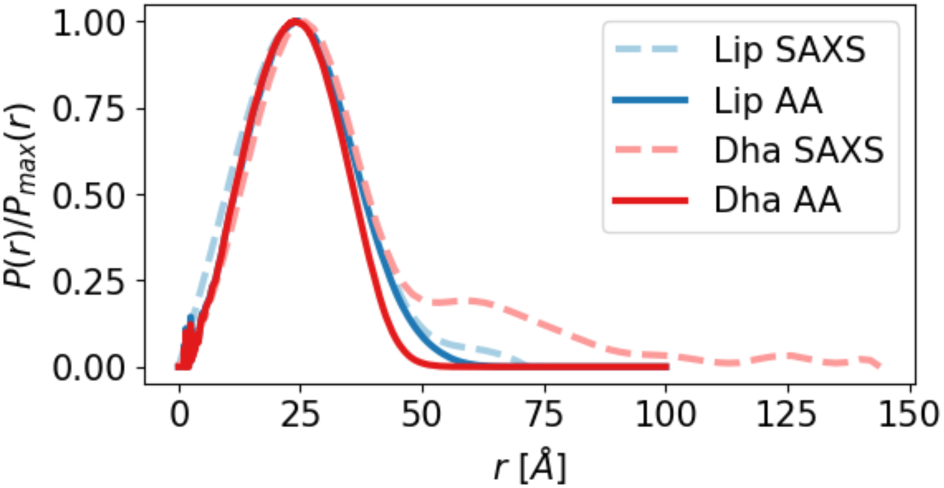
Pair distance distribution function P(r) comparing SAXS measurements and values calculated from all-atom simulation for lipase and dehalogenase.

### 3.2 RMSF - Comparison of Results from Coarse-grained, All-Atom Trajectories and X-ray Diffraction Measurements

Figure 5 shows ensemble average residue-wise RMSF values calculated from all-atom and coarse-grained simulations, along with temperature factor values from the associated PDB structure. There is qualitative agreement between the scales as to which residues along the chain length experience high RMSF values, corresponding to the mobility or flexibility of the protein at that location.

### 3.3 Gyration Tensor - Comparison of Results from All-atom and Coarse-grained Trajectories

Gyration tensor parameter distributions are shown in Figure 4 and summarized in Table 2. In dehalogenase, the sampled ranges for all parameters are very similar between all-atom and coarse-grained. The *R_g_* in both cases matches well with the XRD experiment, indicating that the overall mass distribution of the simulated molecules is reasonable. The *R_g_* distributions are unimodal and highly symmetrical, centered close to 1.8 nm. The width of the main peaks of the all-atom and coarse-grained distributions are also quite similar, suggesting that the structure of the molecule undergoes a similar scale of fluctuation in each model. Values of the anisotropy and asphericity parameters are slightly higher for all-atom trajectories than the coarse-grained trajectories. Such a deviation may be due to a loss in the distinction of characteristic structures within the coarse-grained model owing to its reduced resolution, although values for both the all-atom and coarse-grained models indicate a high degree of sphericity and spherical symmetry.

**Figure 4.**
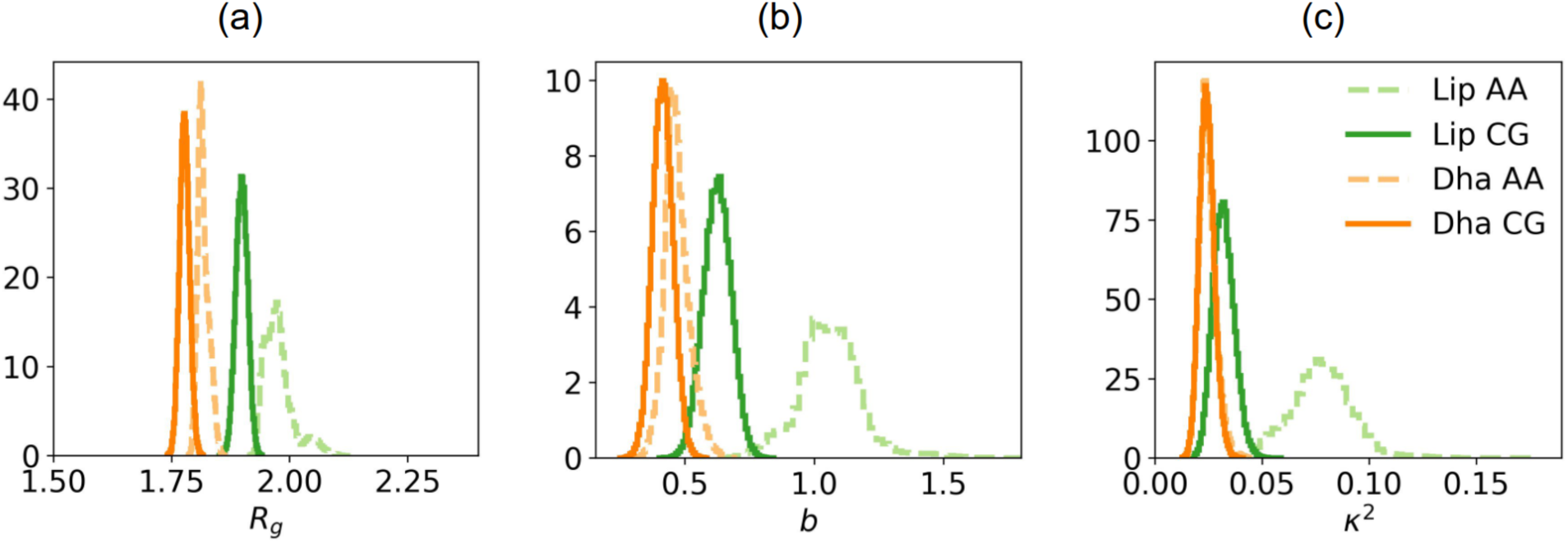
Ensemble distributions of gyration tensor shape parameters. (a) *R_g_*; (b) *b*; (c) *κ^2^*.

**Figure 5.**
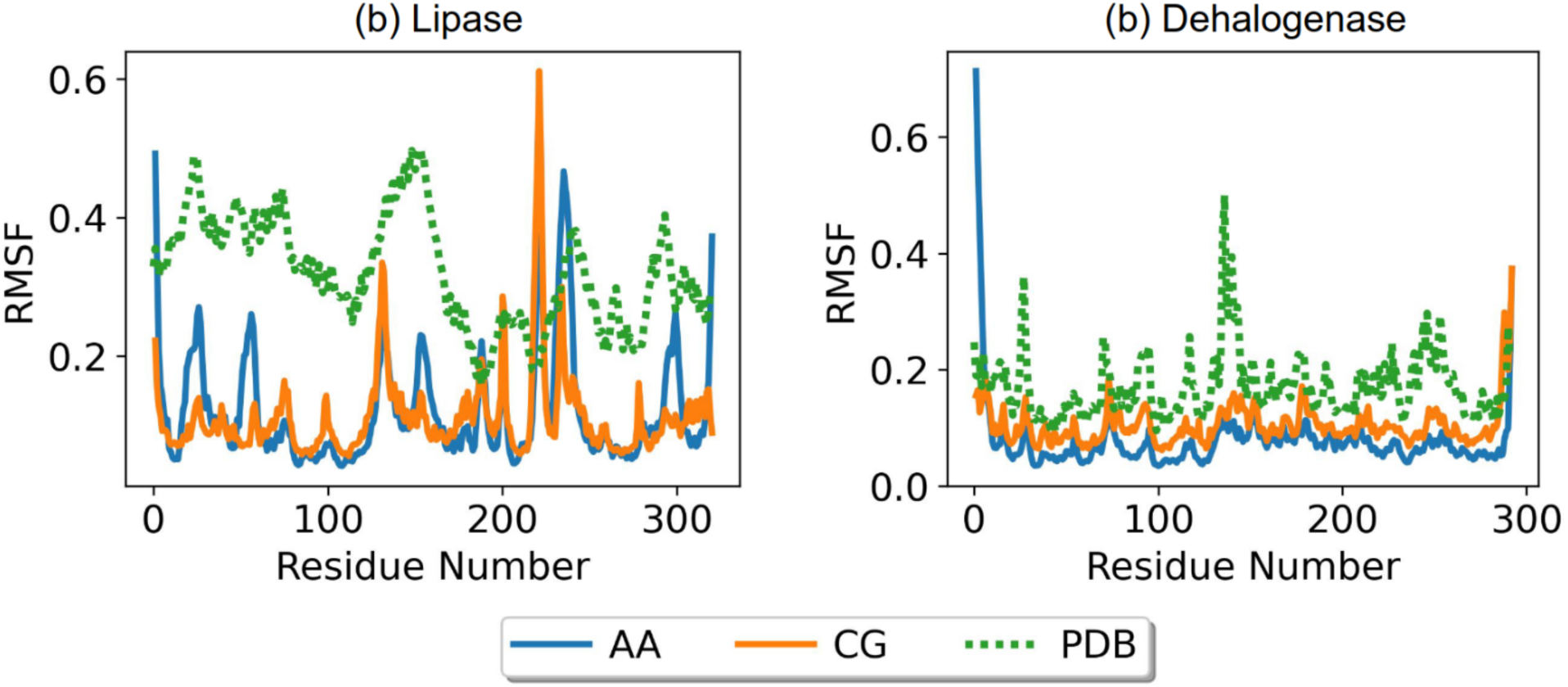
Ensemble average of residue-wise RMSF for all-atom and coarse-grained representations of lipase (left) and dehalogenase (right). Temperature factor data from reference PDB structures are shown as green dotted lines, scaled arbitrarily for visual comparison.

The all-atom model of lipase shows a much broader and asymmetrical distribution of *R_g_*, with the major peak centered around 1.97 nm, compared to 1.89 nm in the coarse-grained model. A minor peak in the all-atom distribution at 2.05 nm represents a small number of configurations which vary significantly from those seen in the majority of the sampled ensemble. As with dehalogenase, the widths of the main peaks in the lipase parameter distributions are similar between all-atom and coarse-grained trajectories. Likewise, the lipase values for anisotropy and asphericity indicate high spherical symmetry.

### 3.4 Surface Chemistry - Comparison of Results from All-Atom and Coarse-grained Trajectories

This study aims to develop a framework for coarse-grained models of enzymes which reproduce their surface chemistry. Therefore, a method is required that quantifies the surface chemistry of enzymes. To this end, the SURFMAP software [Schweke 2022; I2bc/surfmap (Github) 2024] is used to calculate surface characteristics such as electrostatic potential, hydrophobicity, and circular variance of the coarse-grained representations of the enzymes.

#### 3.4.1 Calculated Surface Properties

SURFMAP takes a model of a protein as input and calculates chemical properties on its surface. First, the target property is calculated at the scale of atoms or residues. A point cloud located 3 Å from the surface of the protein is then calculated using the MSMS software [Sanner 1996]. Each point in this surface-approximating cloud takes on the value of the property of interest calculated for the nearest atom in the protein model. The locations of the cloud points are expressed in polar coordinates with the protein center of mass as the origin, creating the basis for a spherical representation. The corresponding property values are then mapped onto a 2D equal-area projection of the spherical surface. This projection uses bins of interval 5 degrees, and the value of the property associated with each bin is the average of the points in the bin.

The surface properties calculated for this study are electrostatic potential, hydrophobicity, and circular variance. SURFMAP utilizes the APBS software [Jurrus 2018] to calculate atom-wise (or pseudoatom-wise) electrostatic potentials, with charge values reported in units of kT/e. For electrostatics at the coarse-grained scale, the user must include a custom PQR file [Dolinsky 2004; PQR molecular structure format 2024] in the input which details pseudoatom configuration and electrostatic charge, which is accomplished using custom codes available in the GitHub repository [Hooten 2024]. Hydrophobicity is calculated in the Kyte-Doolittle scale [Kyte 1982], in which the amino acid residues are assigned index values from -4.5 for most hydrophilic to +4.5 for most hydrophobic. Circular variance (CV) [Mezei 2003] is calculated residue-wise. The CV at a target point is calculated by considering its location relative to the set of points nearby, and may be considered a measure of how near the target point is to the center of the set. CV values range from 0 for exterior points (as in surface protrusions) to 1 for interior points (as in cavities), with the range near 0.5 corresponding to a theoretical surface from which protrusions or cavities extend.

Typical 2D projections showing the surface distribution of electrostatic potential in the all-atom and coarse-grained representations of lipase are shown in Figure 6, illustrating the effect of particle count on the diffuseness of charged surface patches. The scale and resolution of contours in the maps of circular variance and hydrophobicity agree well between all-atom and coarse-grained scales, owing to the fact that these properties are calculated by residue at both scales (additional representative maps of of surface features may be found in the Supporting Information, Figure S3). The hydrophobic maps of all-atom and coarse-grained representations of lipase are largely similar, with values ranging from -4.5 to 4.5 in the Kyte-Doolittle hydrophobicity scale. The data underlying the maps in Figure 6 may be parsed directly to estimate the proportion of the enzyme surface for which properties fall within a range of interest. For example, the section of the surface area that exhibits positive hydrophobicity can be computed as it is a potentially useful property for distinguishing the surface adsorption propensity of a protein.

**Figure 6.**
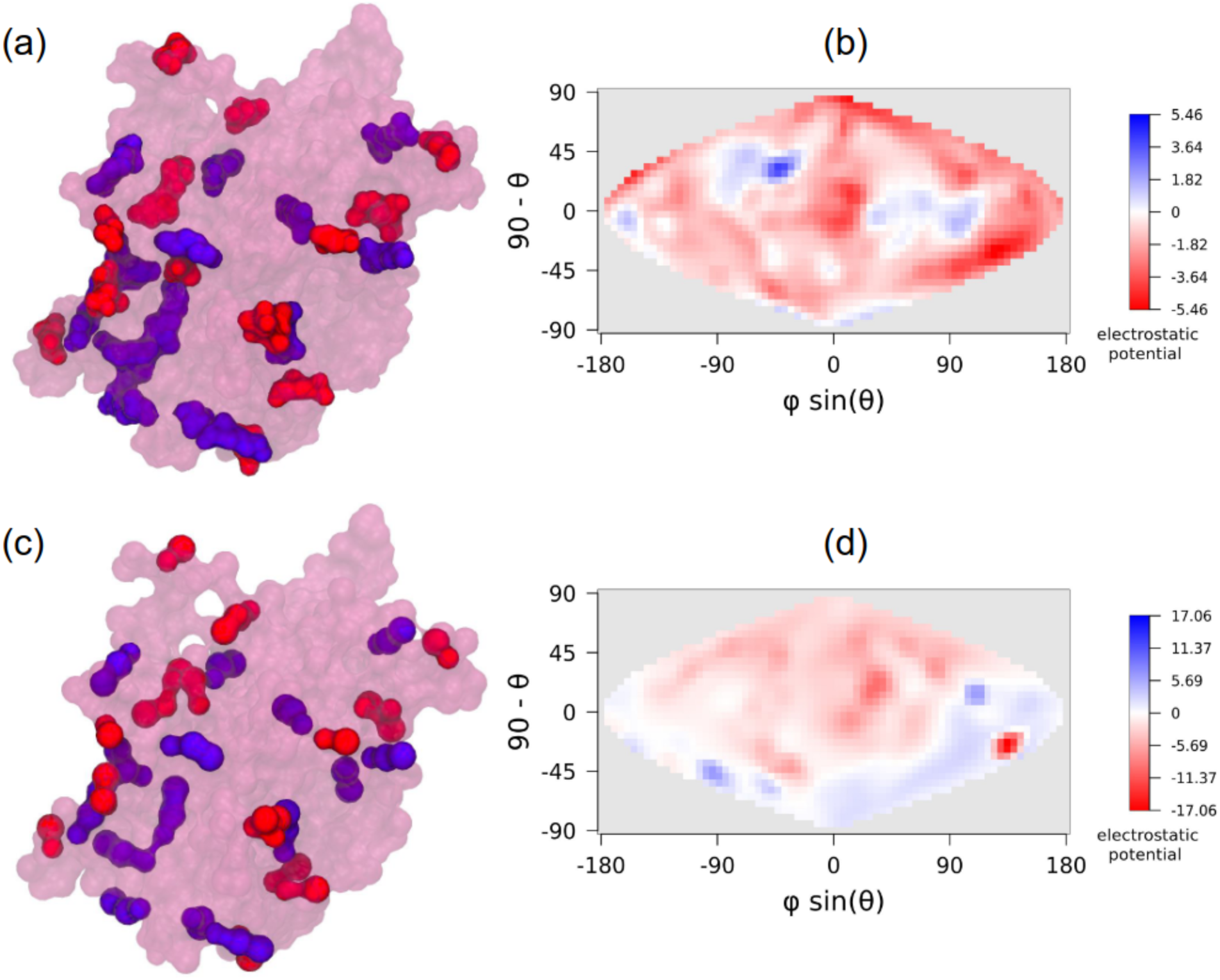
Typical projections of surface distribution of electrostatic potential for all-atom and coarse-grained models of lipase. Panel (a) shows a trajectory frame of the all-atom representation of lipase in which positively charged residues on the surface are labeled in blue, and negatively charged residues in red. Panel (b) shows the associated SURFMAP output for electrostatic potential. Panels (c) and (d) show the same frame as it has been mapped to coarse-grained representation, with its associated SURFMAP calculation

#### 3.4.2 Surface Property Results

Calculated distributions of the surface properties are shown in Figure 7 and summarized in Table 3. The electrostatic potential on the surface of the all-atom representation of lipase ranges from -9 to +5, while the coarse-grained model samples a much larger range from -47 to +26. Similar behavior is observed for dehalogenase, where the potential in the all-atom model ranges from -13 to +7 and in the coarse-grained model from -41 to +26. The areas of the extreme electrostatic potential on the surface of the simulated enzymes are more sharply localized in the coarse-grained representation than in the all-atom representation, as can be seen in Figure 6 and Figure S3. This difference is due to the effective ‘size’ of the coarse-grained beads. The effect is that the bulk of the surface tends more closely to 0 potential, but renders the extreme values as a few small, disjoint spots in contrast to their more gradual distribution in the all-atom representation.

**Figure 7.**
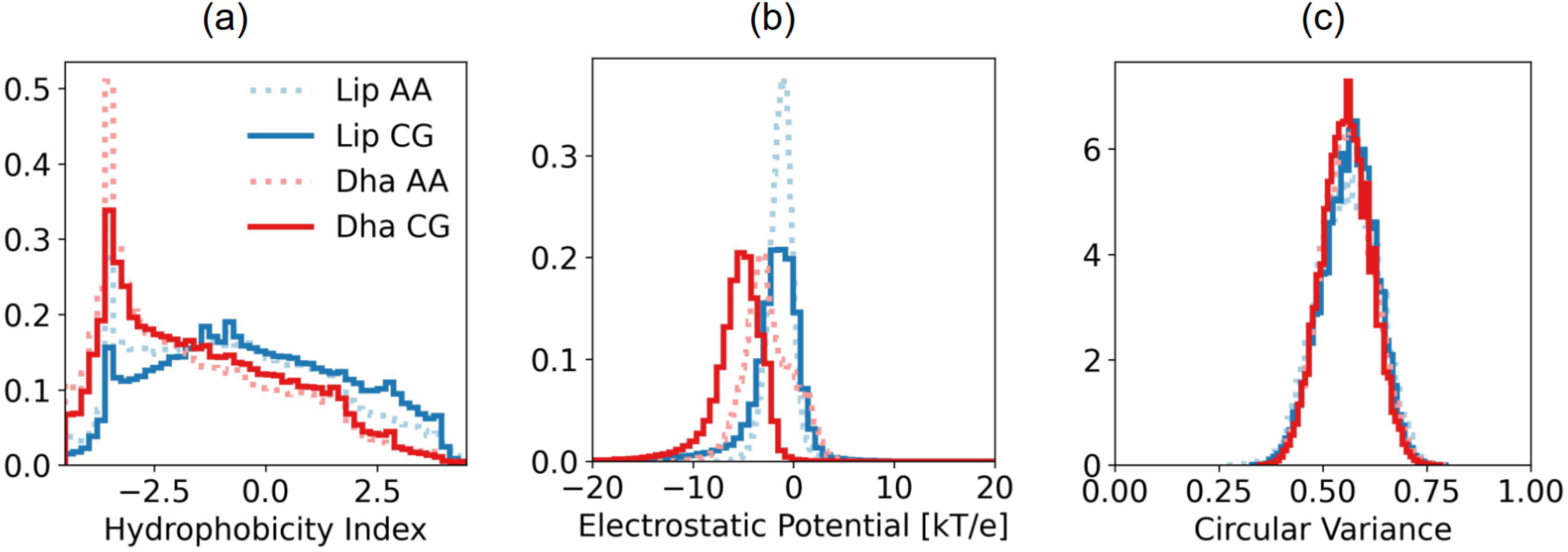
Ensemble histograms of surface property values. (a) Hydrophobicity index ranges from most hydrophilic at -4.5 to most hydrophobic at +4.5. (b) Electrostatic potentials in units of kT/e. (c) Circular variance values range from 0 for points of local surface protrusion to 1 for local concavity.

**Table 3.**
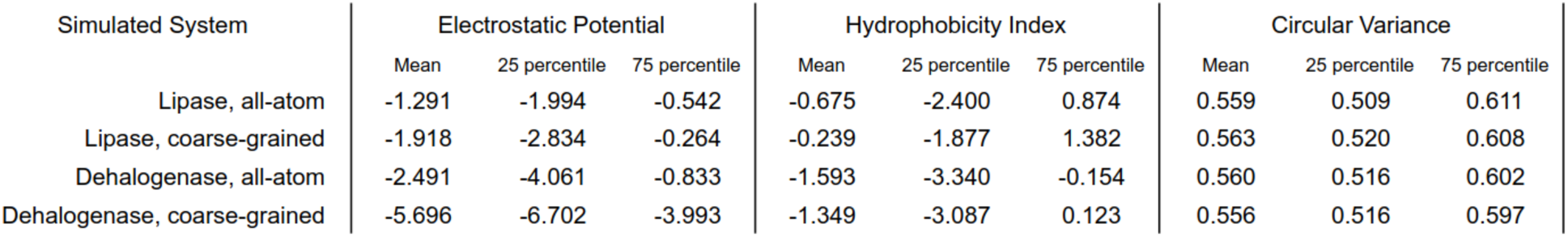
Summary of calculated surface property values.

The distribution of surface potentials in the lipase coarse-grained model deviates slightly from the all-atom reference, showing a slight left skewness. Mean surface potential in lipase is -1.29 in all-atom compared with -1.92 in coarse-grained. In the dehalogenase models the mean is -2.5 in all-atom and -5.7 in coarse-grained.

In all-atom models of both lipase and dehalogenase the hydrophobicity distribution has its highest peak at -3.5, with a much higher peak in dehalogenase than in lipase. This value indicates high hydrophilicity and corresponds in the Kyte-Doolittle scale to aspartic acid, glutamic acid, asparagine, and glutamine. It can be seen in Figure 8 that although ASN and GLN are slightly more common on the surface of lipase than of dehalogenase, dehalogenase has a much greater surface presence of ASP and GLU than lipase. The coarse-grained model of dehalogenase qualitatively matches all-atom almost perfectly, with a slight shift in the mass of the distribution from highly hydrophilic to zero hydrophobicity. The lipase coarse-grained distribution also has an attenuated -3.5 peak compared to all-atom, but in this case, it is associated with an erroneous increase in high hydrophobicity values between 2 and 4. This is very likely due to the higher frequency surface exposure of PHE and VAL in the lipase coarse-grained model compared to its all-atom counterpart, as can be seen in Figure 8.

**Figure 8.**
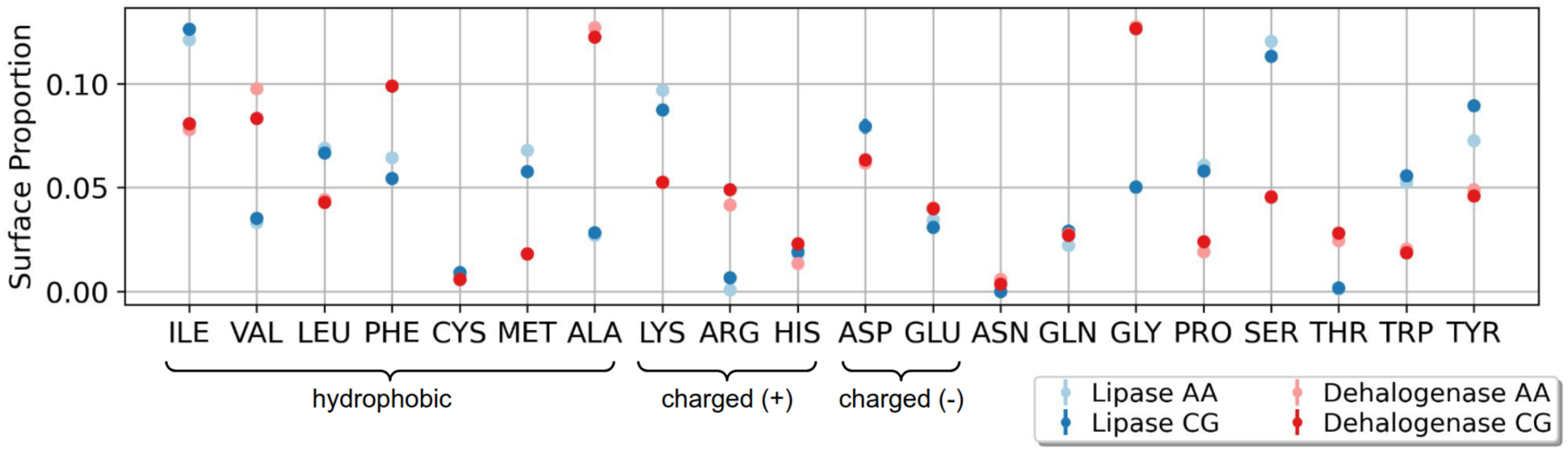
Mean proportion of amino acid residues on the surface of simulated enzymes as a function of amino acids, for all-atom vs coarse-grained representations, with ensemble standard error. Groups are indicated for residues with a positive hydropathy value on the Kyte-Doolittle scale and for the charged residues.

Circular variance distributions are remarkably similar among the two enzyme systems as well as their two modeled resolutions. These distributions are once again unimodal and highly symmetrical with an average value of 0.56 in all four ensembles.

## 4. Conclusions

In this study, two enzyme systems, lipase and dehalogenase, were developed at all-atom and coarse-grained resolution which were validated using SAXS measurements and the structure of corresponding proteins in solution. Values of *R_g_* determined via SAXS fall close to the ranges calculated from simulated MD trajectories, indicating similar mass distributions. In addition, the mode of the experimentally determined PDF agrees well with values from simulations, suggesting the globular and monomeric form of the simulated proteins are in agreement with the experimental sample.

Distributions of the gyration tensor shape parameter values qualitatively agree between the coarse-grained and all-atom models and show a similar scale of fluctuation between resolutions. The calculated distributions are also visually distinctive for the different enzyme systems, a feature likely to be important for future development of the analytical framework.

Likewise, the surfaces calculated using a molecular cartography approach were similar between the all-atom and coarse-grained resolutions, and distinctive between the enzymes. The frequencies at which specific residues occur on the surface of the modeled proteins show that the surface of the coarse-grained model very closely approximates the surface composition of the all-atom reference.

The protocols developed here for the development of coarse-grained models of enzymes and their validation using experimental measurements lay the foundation of a framework which can be used for future efforts in the high-throughput development of such models. These models can be employed to guide experimental efforts towards the stabilization of structure-function relations of protein sequences via stabilization by heteropolymers.

## 5. ACKNOWLEDGEMENTS

This work was funded by the National Science Foundation (DMREF-2118860, DMR-1654325, OAC-1835449 and CBET-2309852). The authors acknowledge the use of computational resources enabled via an allocation from NSF ACCESS (allocation DMR-140125).

MH would like to thank the SURFMAP [Schweke 2022; I2bc/SURFMAP 2024] developers and authors Dr. Anne Lopes (Université Paris-Saclay), Dr. Hugo Schweke (Université Paris-Saclay and Weizmann Institute of Science), and Nicolas Chevrollier for support in the usage of SURFMAP. MH would also like to thank MARTINI developer Fabian Grünewald (University of Groningen) for helpful conversations regarding elastic network model development in Martini.

The authors acknowledge James Byrnes, beamline scientist at NSLS-II beamline 16-ID for Life Science X-ray Scattering (LiX), for his assistance with conducting experiments at Brookhaven National Laboratory. The LiX beamline is part of the Center for BioMolecular Structure (CBMS), which is primarily supported by the National Institutes of Health, National Institute of General Medical Sciences (NIGMS) through a P30 Grant (P30GM133893), and by the DOE Office of Biological and Environmental Research (KP1605010). LiX also received additional support from NIH Grant S10 OD012331. As part of NSLS-II, a national user facility at Brookhaven National Laboratory, work performed at the CBMS was supported in part by the U.S. Department of Energy, Office of Science, Office of Basic Energy Sciences Program under contract number DE-SC0012704. The authors acknowledge Srinivas Chakravartula, Senior Mass Spectrometrist, Wright-Rieman Laboratories, Rutgers University-New Brunswick. His expertise and support greatly contributed to the success of this study.

## References

1. Abraham, M. J.; Murtola, T.; Schulz, R.; Páll, S.; Smith, J. C.; Hess, B.; Lindahl, E. GROMACS: High Performance Molecular Simulations through Multi-Level Parallelism from Laptops to Supercomputers. SoftwareX 2015, 1–2, 19–25. 10.1016/j.softx.2015.06.001.

2. Allen, M. P.; Tildesley, D. J. Computer Simulation of Liquids, Second edition.; Oxford University Press: Oxford, United Kingdom, 2017a.

3. Allen, M. P.; Tildesley, D. J. examples, 2017b. GitHub repository. https://github.com/Allen-Tildesley/examples/blob/master/python_examples/pair_distribution.py. (accessed YYYY-MM-DD)

4. Aydin, F.; Dutt, M. Bioinspired Vesicles Encompassing Two-Tail Phospholipids: Self-Assembly and Phase Segregation via Implicit Solvent Coarse-Grained Molecular Dynamics. J. Phys. Chem. B 2014, 118 (29), 8614–8623. 10.1021/jp503376r.

5. Aydin, F.; Dutt, M. Surface Reconfiguration of Binary Lipid Vesicles via Electrostatically Induced Nanoparticle Adsorption. J. Phys. Chem. B 2016, 120 (27), 6646–6656. 10.1021/acs.jpcb.6b02334.

6. Bahar, I.; Atilgan, A. R.; Erman, B. Direct Evaluation of Thermal Fluctuations in Proteins Using a Single-Parameter Harmonic Potential. Folding and Design 1997, 2 (3), 173–181. 10.1016/S1359-0278(97)00024-2.

7. Banerjee, A.; Hooten, M.; Srouji, N.; Welch, R.; Shovlin, J.; Dutt, M. A Perspective on Coarse-Graining Methodologies for Biomolecules: Resolving Self-Assembly over Extended Spatiotemporal Scales. Front. Soft Matter 2024, 4. 10.3389/frsfm.2024.1361066.

8. Banerjee, A.; Dutt, M. A Hybrid Approach for Coarse-Graining Helical Peptoids: Solvation, Secondary Structure, and Assembly. The Journal of Chemical Physics 2023, 158 (11), 114105. 10.1063/5.0138510.

9. Banerjee, A.; Lu, C. Y.; Dutt, M. A Hybrid Coarse-Grained Model for Structure, Solvation and Assembly of Lipid-like Peptides. Phys. Chem. Chem. Phys. 2022, 24 (3), 1553–1568. 10.1039/D1CP04205J.

10. Berman, H. M.; Westbrook, J.; Feng, Z.; Gilliland, G.; Bhat, T. N.; Weissig, H.; Shindyalov, I. N.; Bourne, P. E. The Protein Data Bank. Nucleic acids research 2000, 28 (1), 235–242. 10.1093/nar/28.1.235

11. Berendsen, H. J. C.; Postma, J. P. M.; van Gunsteren, W. F.; DiNola, A.; Haak, J. R. Molecular Dynamics with Coupling to an External Bath. The Journal of Chemical Physics 1984, 81 (8), 3684–3690. 10.1063/1.448118.

12. Berendsen, H. J. C.; van der Spoel, D.; van Drunen, R. GROMACS: A Message-Passing Parallel Molecular Dynamics Implementation. Computer Physics Communications 1995, 91 (1), 43–56. 10.1016/0010-4655(95)00042-E.

13. Best, R. B.; Zhu, X.; Shim, J.; Lopes, P. E. M.; Mittal, J.; Feig, M.; MacKerell, A. D. Optimization of the Additive CHARMM All-Atom Protein Force Field Targeting Improved Sampling of the Backbone ϕ, ψ and Side-Chain χ _1_ and χ _2_ Dihedral Angles. J. Chem. Theory Comput. 2012, 8 (9), 3257–3273. 10.1021/ct300400x.

14. Bornscheuer, U.; Reif, O.-W.; Lausch, R.; Freitag, R.; Scheper, T.; Kolisis, F. N.; Menge, U. Lipase of Pseudomonas Cepacia for Biotechnological Purposes: Purification, Crystallization and Characterization. Biochimica et Biophysica Acta (BBA) - General Subjects 1994, 1201 (1), 55–60. 10.1016/0304-4165(94)90151-1.

15. Brown, S. T.; Buitrago, P.; Hanna, E.; Sanielevici, S.; Scibek, R.; Nystrom, N. A. Bridges-2: A Platform for Rapidly-Evolving and Data Intensive Research. In Practice and Experience in Advanced Research Computing; PEARC ’21; Association for Computing Machinery: New York, NY, USA, 2021; pp 1–4. 10.1145/3437359.3465593.

16. Bussi, G.; Donadio, D.; Parrinello, M. Canonical Sampling through Velocity Rescaling. The Journal of Chemical Physics 2007, 126 (1), 014101. 10.1063/1.2408420.

17. Carmichael, S. P.; Shell, M. S. A New Multiscale Algorithm and Its Application to Coarse-Grained Peptide Models for Self-Assembly. The Journal of Physical Chemistry B 2012, 116 (29), 8383–8393.

18. Chapman, J.; Ismail, A. E.; Dinu, C. Z. Industrial Applications of Enzymes: Recent Advances, Techniques, and Outlooks. Catalysts 2018, 8 (6), 238. 10.3390/catal8060238.

19. Chapman, R.; Stenzel, M. H. All Wrapped up: Stabilization of Enzymes within Single Enzyme Nanoparticles. J. Am. Chem. Soc. 2019, 141 (7), 2754–2769. 10.1021/jacs.8b10338.

20. Chong, L.; Aydin, F.; Dutt, M. Implicit Solvent Coarse-Grained Model of Polyamidoamine Dendrimers: Role of Generation and pH. Journal of Computational Chemistry 2016, 37 (10), 920–926. 10.1002/jcc.24277.

21. Coupland, C. E.; Andrei, S. A.; Ansell, T. B.; Carrique, L.; Kumar, P.; Sefer, L.; Schwab, R. A.; Byrne, E. F. X.; Pardon, E.; Steyaert, J.; Magee, A. I.; Lanyon-Hogg, T.; Sansom, M. S. P.; Tate, E. W.; Siebold, C. Structure, Mechanism, and Inhibition of Hedgehog Acyltransferase. Molecular Cell 2021, 81 (24), 5025–5038.e10. 10.1016/j.molcel.2021.11.018.

22. Darden, T.; York, D.; Pedersen, L. Particle Mesh Ewald: An N⋅ Log (N) Method for Ewald Sums in Large Systems. The Journal of chemical physics 1993, 98 (12), 10089–10092. 10.1063/1.464397.

23. De Jong, R. M. Structure and Mechanism of a Bacterial Haloalcohol Dehalogenase: A New Variation of the Short-Chain Dehydrogenase/Reductase Fold without an NAD(P)H Binding Site. The EMBO Journal 2003, 22 (19), 4933–4944. 10.1093/emboj/cdg479.

24. Dima, R. I.; Thirumalai, D. Asymmetry in the Shapes of Folded and Denatured States of Proteins. J. Phys. Chem. B 2004, 108 (21), 6564–6570. 10.1021/jp037128y.

25. Dolinsky, T. J.; Nielsen, J. E.; McCammon, J. A.; Baker, N. A. PDB2PQR: An Automated Pipeline for the Setup of Poisson–Boltzmann Electrostatics Calculations. Nucleic Acids Research 2004, 32 (suppl_2), W665–W667. 10.1093/nar/gkh381.

26. Durumeric, A. E. P.; Charron, N. E.; Templeton, C.; Musil, F.; Bonneau, K.; Pasos-Trejo, A. S.; Chen, Y.; Kelkar, A.; Noé, F.; Clementi, C. Machine Learned Coarse-Grained Protein Force-Fields: Are We There Yet? Current Opinion in Structural Biology 2023, 79, 102533. 10.1016/j.sbi.2023.102533.

27. Franke, D.; Petoukhov, M. V.; Konarev, P. V.; Panjkovich, A.; Tuukkanen, A.; Mertens, H. D. T.; Kikhney, A. G.; Hajizadeh, N. R.; Franklin, J. M.; Jeffries, C. M. ATSAS 2.8: A Comprehensive Data Analysis Suite for Small-Angle Scattering from Macromolecular Solutions. Journal of applied crystallography 2017, 50 (4), 1212–1225.

28. Gutiérrez-Fernández, J.; Kaszuba, K.; Minhas, G. S.; Baradaran, R.; Tambalo, M.; Gallagher, D. T.; Sazanov, L. A. Key Role of Quinone in the Mechanism of Respiratory Complex I. Nat Commun 2020, 11 (1), 4135. 10.1038/s41467-020-17957-0.

29. Hedrich, H. C.; Spener, F.; Menge, U.; Hecht, H.-J.; Schmid, R. D. Large-Scale Purification, Enzymic Characterization, and Crystallization of the Lipase from Geotrichum Candidum. Enzyme and microbial technology 1991, 13 (10), 840–847.

30. Hooten, M.; Dutt, M. Enzyme characterization, 2024. GitHub repository. https://github.com/duttm/Polymer_Protein_Hybrid_Particles_DMREF_Team_codes (accessed 2024-09-22).

31. Hooten, M.; Banerjee, A.; Dutt, M. Multiscale, Multiresolution Coarse-Grained Model via a Hybrid Approach: Solvation, Structure, and Self-Assembly of Aromatic Tripeptides. J. Chem. Theory Comput. 2023. 10.1021/acs.jctc.3c00458.

32. Hooten, M.; Dutt, M. Transfer of coarse-grained potentials to new chain lengths in short aromatic peptides [Manuscript in preparation]. Biomedical Engineering, Rutgers University. 2025.

33. Hoover, W. G. Canonical Dynamics: Equilibrium Phase-Space Distributions. Phys. Rev. A 1985, 31 (3), 1695–1697. 10.1103/PhysRevA.31.1695.

34. Hopkins, J. B.; Gillilan, R. E.; Skou, S. BioXTAS RAW: Improvements to a Free Open-Source Program for Small-Angle X-Ray Scattering Data Reduction and Analysis. Journal of applied crystallography 2017, 50 (5), 1545–1553.

35. Hopkins, J. B.; Gillilan, R. E.; Skou, S. BioXTAS RAW: A Free Open-Source Program for Small-Angle X-Ray Scattering Data Reduction. Acta Cryst 2018, 74, a219.

36. Hopkins, J.; Gillilan, R.; Skou, S. BioXTAS RAW 2.0: The Latest in SAXS Data Analysis. Acta Crystallogr A: Found Adv 2020, 76, A27.

37. Huang, J.; Rauscher, S.; Nawrocki, G.; Ran, T.; Feig, M.; De Groot, B. L.; Grubmüller, H.; MacKerell Jr, A. D. CHARMM36m: An Improved Force Field for Folded and Intrinsically Disordered Proteins. Nature methods 2017, 14 (1), 71–73. 10.1038/nmeth.4067

38. I2bc/SURFMAP, 2024. https://github.com/i2bc/SURFMAP (accessed 2024-04-22).

39. Izvekov, S.; Voth, G. A. Multiscale Coarse-Graining of Mixed Phospholipid/Cholesterol Bilayers. J. Chem. Theory Comput. 2006, 2 (3), 637–648. 10.1021/ct050300c.

40. Jorgensen, W. L.; Chandrasekhar, J.; Madura, J. D.; Impey, R. W.; Klein, M. L. Comparison of Simple Potential Functions for Simulating Liquid Water. The Journal of chemical physics 1983, 79 (2), 926–935. 10.1063/1.445869

41. Jurrus, E.; Engel, D.; Star, K.; Monson, K.; Brandi, J.; Felberg, L. E.; Brookes, D. H.; Wilson, L.; Chen, J.; Liles, K.; Chun, M.; Li, P.; Gohara, D. W.; Dolinsky, T.; Konecny, R.; Koes, D. R.; Nielsen, J. E.; Head-Gordon, T.; Geng, W.; Krasny, R.; Wei, G.-W.; Holst, M. J.; McCammon, J. A.; Baker, N. A. Improvements to the APBS Biomolecular Solvation Software Suite. Protein Science 2018, 27 (1), 112–128. 10.1002/pro.3280.

42. Keefe, A. J.; Jiang, S. Poly(Zwitterionic)Protein Conjugates Offer Increased Stability without Sacrificing Binding Affinity or Bioactivity. Nature Chem 2012, 4 (1), 59–63. 10.1038/nchem.1213.

43. Kim, K. K.; Song, H. K.; Shin, D. H.; Hwang, K. Y.; Suh, S. W. The Crystal Structure of a Triacylglycerol Lipase from Pseudomonas Cepacia Reveals a Highly Open Conformation in the Absence of a Bound Inhibitor. Structure 1997, 5 (2), 173–185. 10.1016/S0969-2126(97)00177-9.

44. Ko, J. H.; Maynard, H. D. A Guide to Maximizing the Therapeutic Potential of Protein– Polymer Conjugates by Rational Design. Chemical Society Reviews 2018, 47 (24), 8998–9014. 10.1039/C8CS00606G.

45. Kosuri, S.; Borca, C. H.; Mugnier, H.; Tamasi, M.; Patel, R. A.; Perez, I.; Kumar, S.; Finkel, Z.; Schloss, R.; Cai, L.; Yarmush, M. L.; Webb, M. A.; Gormley, A. J. Machine-Assisted Discovery of Chondroitinase ABC Complexes toward Sustained Neural Regeneration. Advanced Healthcare Materials 2022, 11 (10), 2102101. 10.1002/adhm.202102101.

46. Kroon, P. C.; Grünewald, F.; Barnoud, J.; van Tilburg, M.; Souza, P. C. T.; Wassenaar, T. A.; Marrink, S.-J. Martinize2 and Vermouth: Unified Framework for Topology Generation. arXiv June 30, 2023. 10.48550/arXiv.2212.01191.

47. Kyte, J.; Doolittle, R. F. A Simple Method for Displaying the Hydropathic Character of a Protein. Journal of Molecular Biology 1982, 157 (1), 105–132. 10.1016/0022-2836(82)90515-0.

48. Lahoda, M.; Mesters, J. R.; Stsiapanava, A.; Chaloupkova, R.; Kuty, M.; Damborsky, J.; Kuta Smatanova, I. Crystallographic Analysis of 1,2,3-Trichloropropane Biodegradation by the Haloalkane Dehalogenase DhaA31. Acta Cryst D 2014, 70 (2), 209–217. 10.1107/S1399004713026254.

49. Lancaster, L.; Abdallah, W.; Banta, S.; Wheeldon, I. Engineering Enzyme Microenvironments for Enhanced Biocatalysis. Chem. Soc. Rev. 2018, 47 (14), 5177–5186. 10.1039/c8cs00085a.

50. Li, J.; Jin, K.; C. Mushnoori, S.; Dutt, M. Mechanisms Underlying Interactions between PAMAM Dendron-Grafted Surfaces with DPPC Membranes. RSC Advances 2018, 8 (44), 24982–24992. 10.1039/C8RA03742F.

51. Lindahl, E.; Hess, B.; van der Spoel, D. GROMACS 3.0: A Package for Molecular Simulation and Trajectory Analysis. J Mol Model 2001, 7 (8), 306–317. 10.1007/s008940100045.

52. Luić, M.; Tomić, S.; Leščić, I.; Ljubović, E.; Šepac, D.; Šunjić, V.; Vitale, L.; Saenger, W.; Kojić-Prodić, B. Complex of Burkholderia Cepacia Lipase with Transition State Analogue of 1-phenoxy-2-acetoxybutane: Biocatalytic, Structural and Modelling Study. European Journal of Biochemistry 2001, 268 (14), 3964–3973. 10.1046/j.1432-1327.2001.02303.x.

53. Lyubartsev, A. P.; Laaksonen, A. Calculation of Effective Interaction Potentials from Radial Distribution Functions: A Reverse Monte Carlo Approach. Phys. Rev. E 1995, 52 (4), 3730–3737. 10.1103/PhysRevE.52.3730.

54. Manalastas-Cantos, K.; Konarev, P. V.; Hajizadeh, N. R.; Kikhney, A. G.; Petoukhov, M. V.; Molodenskiy, D. S.; Panjkovich, A.; Mertens, H. D.; Gruzinov, A.; Borges, C. ATSAS 3.0: Expanded Functionality and New Tools for Small-Angle Scattering Data Analysis. Journal of applied crystallography 2021, 54 (1), 343–355.

55. Markova, K.; Chmelova, K.; M. Marques, S.; Carpentier, P.; Bednar, D.; Damborsky, J.; Marek, M. Decoding the Intricate Network of Molecular Interactions of a Hyperstable Engineered Biocatalyst. Chemical Science 2020, 11 (41), 11162–11178. 10.1039/D0SC03367G.

56. Markova, K.; Kunka, A.; Chmelova, K.; Havlasek, M.; Babkova, P.; Marques, S. M.; Vasina, M.; Planas-Iglesias, J.; Chaloupkova, R.; Bednar, D.; Prokop, Z.; Damborsky, J.; Marek, M. Computational Enzyme Stabilization Can Affect Folding Energy Landscapes and Lead to Catalytically Enhanced Domain-Swapped Dimers. ACS Catal. 2021, 11 (21), 12864–12885. 10.1021/acscatal.1c03343.

57. Marrink, S. J.; Risselada, H. J.; Yefimov, S.; Tieleman, D. P.; de Vries, A. H. The MARTINI Force Field: Coarse Grained Model for Biomolecular Simulations. The journal of physical chemistry. B 2007, 111 (27), 7812–7824. 10.1021/jp071097f.

58. Marrink, S. J.; Corradi, V.; Souza, P. C. T.; Ingólfsson, H. I.; Tieleman, D. P.; Sansom, M. S. P. Computational Modeling of Realistic Cell Membranes. Chem. Rev. 2019, 119 (9), 6184–6226. 10.1021/acs.chemrev.8b00460.

59. Mezei, M. A New Method for Mapping Macromolecular Topography. Journal of Molecular Graphics and Modelling 2003, 21 (5), 463–472. 10.1016/S1093-3263(02)00203-6.

60. Mushnoori, S.; Lu, C. Y.; Schmidt, K.; Dutt, M. A Coarse-Grained Molecular Dynamics Study of Phase Behavior in Co-Assembled Lipomimetic Oligopeptides. Journal of Molecular Graphics and Modelling 2023, 125, 108624. 10.1016/j.jmgm.2023.108624.

61. Mushnoori, S.; Schmidt, K.; Nanda, V.; Dutt, M. Designing Phenylalanine-Based Hybrid Biological Materials: Controlling Morphology via Molecular Composition. Organic & Biomolecular Chemistry 2018, 16 (14), 2499–2507. 10.1039/C8OB00130H.

62. Muthukumar, V. C.; Chong, L.; Dutt, M. Designing Soft Nanomaterials via the Self Assembly of Functionalized Icosahedral Viral Capsid Nanoparticles. Journal of Materials Research 2015, 30 (1), 141–150. 10.1557/jmr.2014.346.

63. Nardini, M.; Dijkstra, B. W. α/β Hydrolase Fold Enzymes: The Family Keeps Growing. Current Opinion in Structural Biology 1999, 9 (6), 732–737. 10.1016/S0959-440X(99)00037-8.

64. Nosé, S. A Molecular Dynamics Method for Simulations in the Canonical Ensemble. Molecular Physics 1984, 52 (2), 255–268. 10.1080/00268978400101201.

65. Panganiban, B.; Qiao, B.; Jiang, T.; DelRe, C.; Obadia, M. M.; Nguyen, T. D.; Smith, A. A. A.; Hall, A.; Sit, I.; Crosby, M. G.; Dennis, P. B.; Drockenmuller, E.; Olvera de la Cruz, M.; Xu, T. Random Heteropolymers Preserve Protein Function in Foreign Environments. Science (American Association for the Advancement of Science) 2018, 359 (6381), 1239–1243. 10.1126/science.aao0335.

66. Parrinello, M.; Rahman, A. Polymorphic Transitions in Single Crystals: A New Molecular Dynamics Method. Journal of Applied physics 1981, 52 (12), 7182–7190.

67. Pelegri-O’Day, E. M.; Lin, E.-W.; Maynard, H. D. Therapeutic Protein–Polymer Conjugates: Advancing Beyond PEGylation. J. Am. Chem. Soc. 2014, 136 (41), 14323–14332. 10.1021/ja504390x.

68. Peng, Y.; Pak, A. J.; Durumeric, A. E. P.; Sahrmann, P. G.; Mani, S.; Jin, J.; Loose, T. D.; Beiter, J.; Voth, G. A. OpenMSCG: A Software Tool for Bottom-Up Coarse-Graining. J. Phys. Chem. B 2023, 127 (40), 8537–8550. 10.1021/acs.jpcb.3c04473.

69. Pezeshkian, W.; Grünewald, F.; Narykov, O.; Lu, S.; Arkhipova, V.; Solodovnikov, A.; Wassenaar, T. A.; Marrink, S. J.; Korkin, D. Molecular Architecture and Dynamics of SARS-CoV-2 Envelope by Integrative Modeling. Structure 2023, 31 (4), 492–503.e7. 10.1016/j.str.2023.02.006.

70. Poma, A. B.; Cieplak, M.; Theodorakis, P. E. Combining the MARTINI and Structure-Based Coarse-Grained Approaches for the Molecular Dynamics Studies of Conformational Transitions in Proteins. J. Chem. Theory Comput. 2017, 13 (3), 1366–1374. 10.1021/acs.jctc.6b00986.

71. PQR molecular structure format — pdb2pqr 3.6.2 documentation. https://pdb2pqr.readthedocs.io/en/latest/formats/pqr.html (accessed 2024-04-22).

72. RCSB Protein Data Bank. RCSB PDB - 4KAF: Crystal Structure of Haloalkane dehalogenase HaloTag7 at the resolution 1.5A, Northeast Structural Genomics Consortium (NESG) Target OR151. https://www.rcsb.org/structure/4kaf (accessed 2024-09-08).

73. Reddy, T.; Sansom, M. S. P. The Role of the Membrane in the Structure and Biophysical Robustness of the Dengue Virion Envelope. Structure 2016, 24 (3), 375–382. 10.1016/j.str.2015.12.011.

74. Reith, D.; Pütz, M.; Müller-Plathe, F. Deriving Effective Mesoscale Potentials from Atomistic Simulations. Journal of Computational Chemistry 2003, 24 (13), 1624–1636. 10.1002/jcc.10307.

75. Ridder, I. S.; Rozeboom, H. J.; Kalk, K. H.; Dijkstra, B. W. Crystal Structures of Intermediates in the Dehalogenation of Haloalkanoates by L-2-Haloacid Dehalogenase. Journal of Biological Chemistry 1999, 274 (43), 30672–30678.

76. Rios, N. S.; Pinheiro, B. B.; Pinheiro, M. P.; Bezerra, R. M.; dos Santos, J. C. S.; Gonçalves, L. R. B. Biotechnological Potential of Lipases from Pseudomonas: Sources, Properties and Applications. Process biochemistry 2018, 75, 99–120.

77. Ruff, K. M.; Harmon, T. S.; Pappu, R. V. CAMELOT: A Machine Learning Approach for Coarse-Grained Simulations of Aggregation of Block-Copolymeric Protein Sequences. J. Chem. Phys. 2015, 143 (24). 10.1063/1.4935066.

78. Rühle, V.; Junghans, C. Hybrid Approaches to Coarse-Graining Using the VOTCA Package: Liquid Hexane. Macromolecular Theory and Simulations 2011, 20 (7), 472–477. 10.1002/mats.201100011.

79. Sahrmann, P. G.; Loose, T. D.; Durumeric, A. E. P.; Voth, G. A. Utilizing Machine Learning to Greatly Expand the Range and Accuracy of Bottom-Up Coarse-Grained Models through Virtual Particles. J. Chem. Theory Comput. 2023, 19 (14), 4402–4413. 10.1021/acs.jctc.2c01183.

80. Sanner, M. F.; Olson, A. J.; Spehner, J.-C. Reduced Surface: An Efficient Way to Compute Molecular Surfaces. Biopolymers 1996, 38 (3), 305–320. 10.1002/(SICI)1097-0282(199603)38:3<305::AID-BIP4>3.0.CO;2-Y.

81. Shell, M. S. The Relative Entropy Is Fundamental to Multiscale and Inverse Thermodynamic Problems. The Journal of Chemical Physics 2008, 129 (14), 144108. 10.1063/1.2992060.

82. Schweke, H.; Mucchielli, M.-H.; Chevrollier, N.; Gosset, S.; Lopes, A. SURFMAP: A Software for Mapping in Two Dimensions Protein Surface Features. J. Chem. Inf. Model. 2022, 62 (7), 1595–1601. 10.1021/acs.jcim.1c01269.

83. Sirugue, L.; Langenfeld, F.; Lagarde, N.; Montes, M. PLO3S: Protein LOcal Surficial Similarity Screening. Computational and Structural Biotechnology Journal 2024, 26, 1–10. 10.1016/j.csbj.2023.12.002.

84. Souza, P. C. T.; Thallmair, S.; Conflitti, P.; Ramírez-Palacios, C.; Alessandri, R.; Raniolo, S.; Limongelli, V.; Marrink, S. J. Protein–Ligand Binding with the Coarse-Grained Martini Model. Nat Commun 2020, 11 (1), 3714. 10.1038/s41467-020-17437-5.

85. Souza, P. C. T.; Alessandri, R.; Barnoud, J.; Thallmair, S.; Faustino, I.; Grünewald, F.; Patmanidis, I.; Abdizadeh, H.; Bruininks, B. M. H.; Wassenaar, T. A.; Kroon, P. C.; Melcr, J.; Nieto, V.; Corradi, V.; Khan, H. M.; Domański, J.; Javanainen, M.; Martinez-Seara, H.; Reuter, N.; Best, R. B.; Vattulainen, I.; Monticelli, L.; Periole, X.; Tieleman, D. P.; de Vries, A. H.; Marrink, S. J. Martini 3: A General Purpose Force Field for Coarse-Grained Molecular Dynamics. Nat Methods 2021, 18 (4), 382–388. 10.1038/s41592-021-01098-3.

86. Svergun, D. I. Restoring Low Resolution Structure of Biological Macromolecules from Solution Scattering Using Simulated Annealing. Biophysical journal 1999, 76 (6), 2879–2886.

87. Synek, L.; Pleskot, R.; Sekereš, J.; Serrano, N.; Vukašinović, N.; Ortmannová, J.; Klejchová, M.; Pejchar, P.; Batystová, K.; Gutkowska, M.; Janková-Drdová, E.; Marković, V.; Pečenková, T.; Šantrůček, J.; Žárský, V.; Potocký, M. Plasma Membrane Phospholipid Signature Recruits the Plant Exocyst Complex via the EXO70A1 Subunit. Proc. Natl. Acad. Sci. U.S.A. 2021, 118 (36), e2105287118. 10.1073/pnas.2105287118.

88. Tamasi, M. J.; Patel, R. A.; Borca, C. H.; Kosuri, S.; Mugnier, H.; Upadhya, R.; Murthy, N. S.; Webb, M. A.; Gormley, A. J. Machine Learning on a Robotic Platform for the Design of Polymer–Protein Hybrids. Advanced Materials 2022, 2201809. 10.1002/adma.202201809.

89. Theodorou, D. N.; Suter, U. W. Shape of Unperturbed Linear Polymers: Polypropylene. Macromolecules 1985, 18 (6), 1206–1214. 10.1021/ma00148a028.

90. Tirion, M. M. Large Amplitude Elastic Motions in Proteins from a Single-Parameter, Atomic Analysis. Phys. Rev. Lett. 1996, 77 (9), 1905–1908. 10.1103/PhysRevLett.77.1905.

91. Togashi, Y.; Flechsig, H. Coarse-Grained Protein Dynamics Studies Using Elastic Network Models. International Journal of Molecular Sciences 2018, 19 (12), 3899. 10.3390/ijms19123899.

92. Towns, J.; Cockerill, T.; Dahan, M.; Foster, I.; Gaither, K.; Grimshaw, A.; Hazlewood, V.; Lathrop, S.; Lifka, D.; Peterson, G. D.; Roskies, R.; Scott, J. R.; Wilkins-Diehr, N. XSEDE: Accelerating Scientific Discovery. Computing in Science & Engineering 2014, 16 (5), 62–74. 10.1109/MCSE.2014.80.

93. Van Der Spoel, D.; Lindahl, E.; Hess, B.; Groenhof, G.; Mark, A. E.; Berendsen, H. J. C. GROMACS: Fast, Flexible, and Free. Journal of Computational Chemistry 2005, 26 (16), 1701–1718. 10.1002/jcc.20291.

94. Villa, A.; Vegt, N. F. A. van der; Peter, C. Self-Assembling Dipeptides: Including Solvent Degrees of Freedom in a Coarse-Grained Model. Phys. Chem. Chem. Phys. 2009, 11 (12), 2068–2076. 10.1039/B818146M.

95. Volkov, V. V.; Svergun, D. I. Uniqueness of Ab Initio Shape Determination in Small-Angle Scattering. Journal of applied crystallography 2003, 36 (3), 860–864.

96. Wang, Q.; Cheung, M. S. A Physics-Based Approach of Coarse-Graining the Cytoplasm of Escherichia Coli (CGCYTO). Biophysical Journal 2012, 102 (10), 2353–2361. 10.1016/j.bpj.2012.04.010.

97. Wang, B.; Zhong, C.; Tieleman, D. P. Supramolecular Organization of SARS-CoV and SARS-CoV-2 Virions Revealed by Coarse-Grained Models of Intact Virus Envelopes. J. Chem. Inf. Model. 2022, 62 (1), 176–186. 10.1021/acs.jcim.1c01240.

