## Supporting Information File for "Framework for Martini-based Coarse-grained Model of Enzymes: Model Development and Experimental Validation"

### Table S1. Simulated system details

| Enzyme | Scale | Number of Waters | Number of Protein Atoms | Number of Ions | Total Atoms | Mass of modeled protein [amu] |
|---|---|---|---|---|---|---|
| Lipase | AA | 30481 | 4634 | 1 Ca2+, 2 Na+ | 96080 | 33125.0098 |
| | CG | 6878 | 700 | 1 Ca2+, 2 Na+ | 7581 | 38898 |
| Dehalogenase | AA | 23166 | 4644 | 18 Na+, 1 Cl- | 74161 | 33282.0548 |
| | CG | 5105 | 711 | 18 Na+, 1 Cl- | 5385 | 49500 |

### Table S2. Simulation parameters

Tables below show simulation parameters used for AA and CG simulations as indicated.

| AA | | |
|---|---|---|
| Minimization | Max Steps | 50000 |
| | Restraints | NA |
| | Minimization Algorithm | Steepest descent |
| | Force Tolerance | 1000 kJ/mol/nm |
| Equilibration | Integrator | leap-frog |
| | Steps | 500000 |
| | Time Step | 2 fs |
| | Restraints | Harmonic restraints on protein heavy atoms |
| | Constraints | LINCS on bonds with hydrogen |
| | Thermostat | velocity rescaling with a stochastic term |
| | Ref Temperature | 300 K |
| | Barostat | NA |
| | Ref Pressure | NA |
| Production | Integrator | leap-frog |
| | Steps | 250000000 |
| | Time Step | 2 fs |
| | Sampling Interval | 1000 steps |
| | Thermostat | Nose-Hoover |
| | Ref Temperature | 300 K |
| | Barostat | Parrinello-Rahman |
| | Ref Pressure | 1 bar |
| | Electrostatics Evaluation | Particle Mesh Ewald |
| | ES Real Space Cutoff | 1 nm |
| | VDW Evaluation | cutoff |
| | VDW Cutoff Scheme | potential shift |
| | VDW Cutoff Distance | 1 nm |

| CG | | |
|---|---|---|
| Minimization | Max Steps | 50000 |
| | Restraints | NA |
| | Minimization Algorithm | Steepest descent |
| | Force Tolerance | 1000 kJ/mol/nm |
| Equilibration 1 | Integrator | leap-frog |
| | Steps | 2000 |
| | Time Step | 5 fs |
| | Restraints | Harmonic restraints on protein backbone beads |
| | Constraints | NA |
| | Thermostat | velocity rescaling with a stochastic term |
| | Ref Temperature | 300 K |
| | Barostat | NA |
| | Ref Pressure | NA |
| Equilibration 2 | Integrator | leap-frog |
| | Steps | 25000 |
| | Time Step | 2 fs |
| | Restraints | NA |
| | Constraints | NA |
| | Thermostat | velocity rescaling with a stochastic term |
| | Ref Temperature | 300 K |
| | Barostat | NA |
| | Ref Pressure | NA |
| Equilibration 3 | Integrator | leap-frog |
| | Steps | 10000 |
| | Time Step | 10 fs |
| | Restraints | NA |
| | Constraints | NA |
| | Thermostat | velocity rescaling with a stochastic term |
| | Ref Temperature | 300 K |
| | Barostat | Berendsen |

| | | |
|---|---|---|
| | Ref Pressure | 1 bar |
| Production | Integrator | leap-frog |
| | Steps | 25000000 |
| | Time Step | 20 fs |
| | Sampling Interval | 100 steps |
| | Thermostat | Nose-Hoover |
| | Ref Temperature | 300 K |
| | Barostat | Parrinello-Rahman |
| | Ref Pressure | 1 bar |
| | Electrostatics Evaluation | Particle Mesh Ewald |
| | ES Real Space Cutoff | 1 nm |
| | VDW Evaluation | cutoff |
| | VDW Cutoff Scheme | potential shift |
| | VDW Cutoff Distance | 1 nm |

### Feature distributions

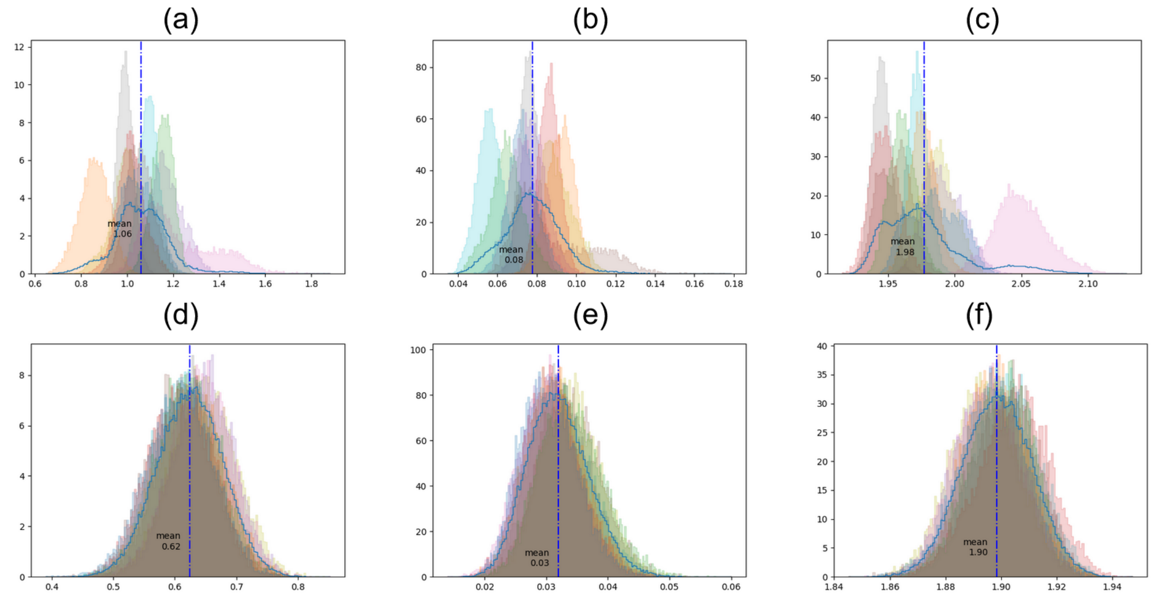

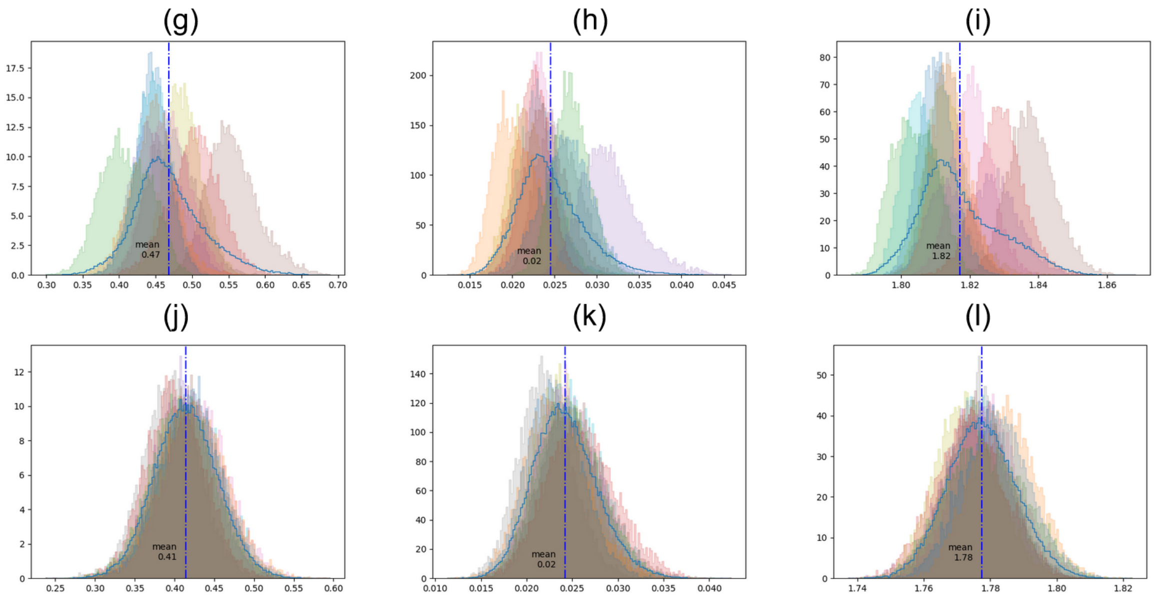


Figure S1. Distributions of gyration tensor shape parameters. Left column asphericity, center column shape anisotropy, right column radius of gyration. Rows from top are: lipase AA (a-c), lipase CG (d-f), dehalogenase AA (g-i), dehalogenase CG (j-l). Individual distributions for ten independent simulations of each system shown shaded. Cumulative distributions for completely pooled data shown with solid blue lines. The AA distributions take on slightly higher values than their CG counterparts, and are more skewed and generally less orderly.

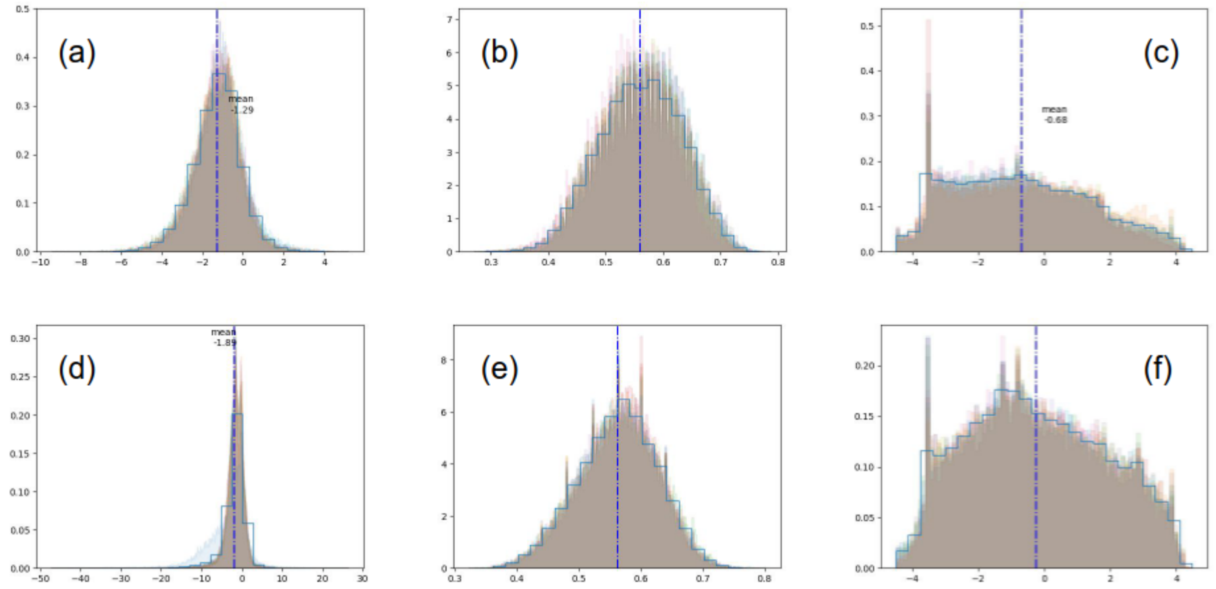

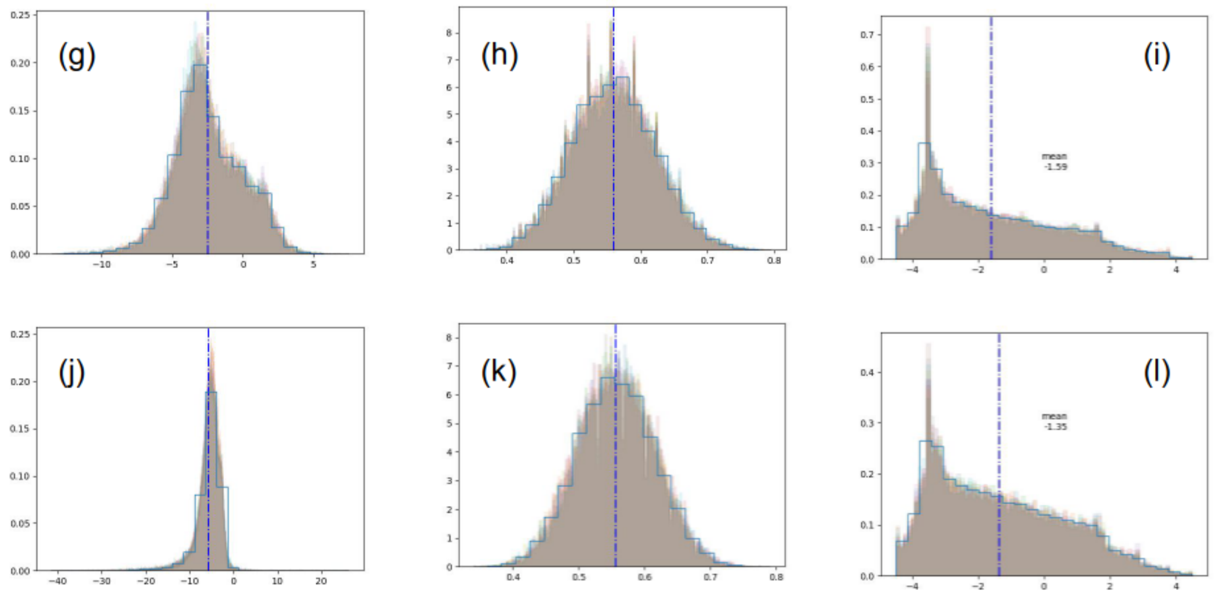


Figure S2. Distributions of calculated surface features. Left column surface electrostatic potential, center column circular variance, right column hydrophobicity. Rows from top are: lipase AA (a-c), lipase CG (d-f), dehalogenase AA (g-i), dehalogenase CG (j-l). Individual distributions for ten independent simulations of each system shown shaded. Cumulative distributions for completely pooled data shown with solid blue lines.

Typical Surface Calculations

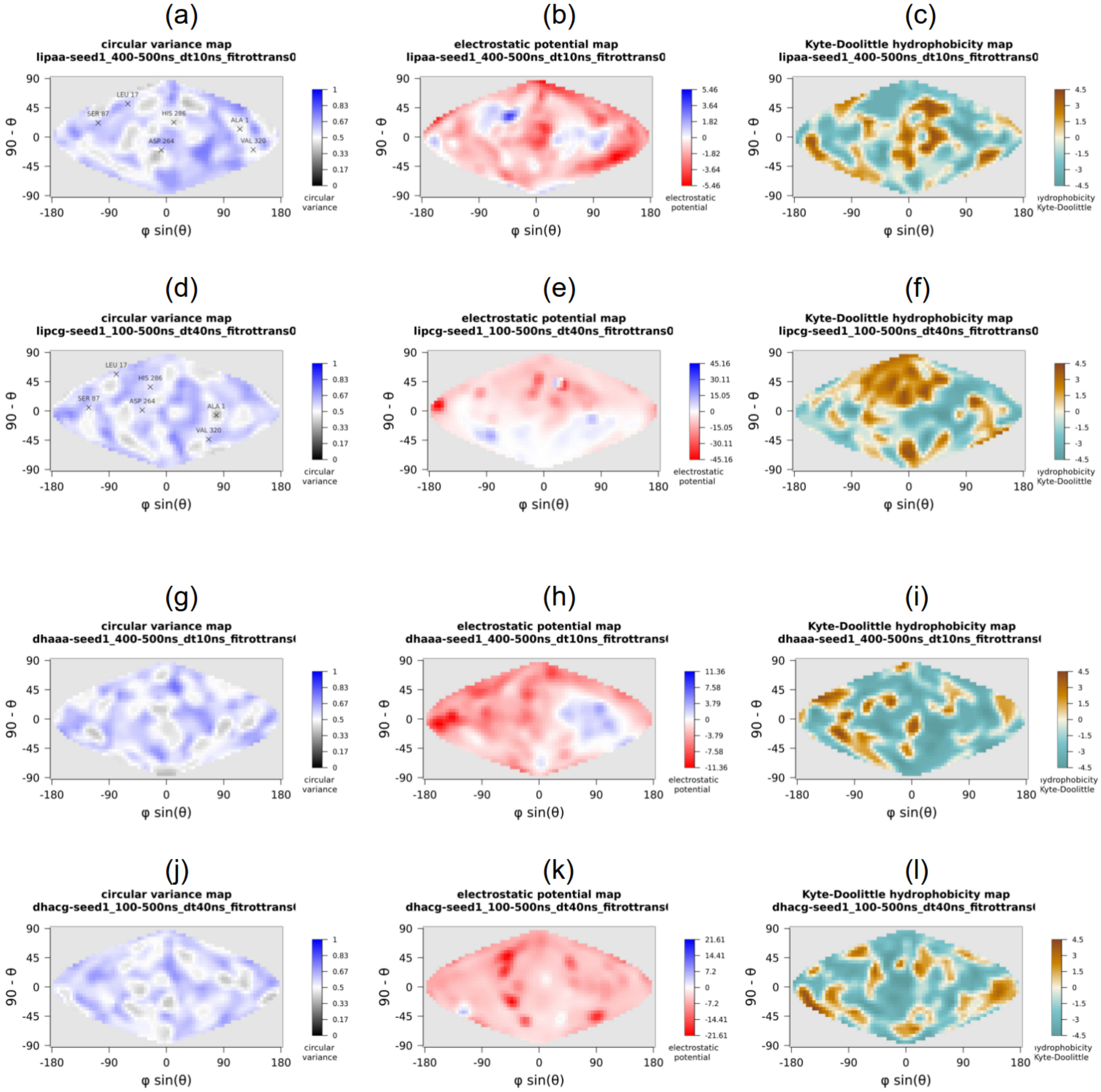


Figure S3. Surface feature values calculated from a typical simulation frame. Left column circular variance, center column electrostatic potential, center, right column hydrophobicity. Rows from top are: lipase AA (a-c), lipase CG (d-f), dehalogenase AA (g-i), dehalogenase CG (j-l).

### Mass spectrometry Experiment details

**Mass Spectrometry and Liquid-Chromatography Mass Spectrometry (LC-MS)**

High resolution mass spectra was obtained on a system, Matrix assisted laser desorption/ionization time of flight mass spectrometry (MALDI-TOF-MS). Data analysis was conducted using Thermo Scientific Xcalibur software.

Sample was prepared desalting with a PD10 column and eluted with a suitable buffer (50mM Ammonium Acetate). The solution was injected into the and separated on an Acquity UPLC BEH C18 column using a gradient elution of water and acetonitrile, containing 0.1% formic acid. The mass spectrometer operated with a spray voltage of 3.5 kV, capillary temperature of 320°C, and S-Lens RF level of 50. Full MS scans were acquired at a resolution of 70,000.

Samples were initially analyzed by LC-MS/MS using a RSLC system (Dionex Ultimate 3000,Dionex, Sunnyvale, CA) interfaced with Velos LTQ –Orbitrap (Thermo-fisher, San Jose, CA). Instrument started with 15% Buffer B (A: 0.2% formic acid in water, B: 0.1% formic acid in acetonitrile) for 3 min with a flow rate of 200 µL/min. The samples were desalted through a reverse phase column- Halo Protein C4 Column (2.1 mm × 50 mm, 3.4 um) using a linear gradient 15-100% B in 7 min, and kept at 100% B for 3 min before equilibrating for the next run. Mass spectrometry data was acquired from the ion trap using positive mode with scan range of m/z 300–2000.For direct infusion, a sample solubilized in 10 mM ammonium ace-tate, pH 8.5 with protein concentration of 4.4 mg/ml was infused to ion source at 8 ul/min. Data was collected at full scan positive ion trap with m/z range of 300–2000. The raw data files were analyzed using Thermo Scientific’s Protein Deconvolution 4.0. Peaks were manually deconvosluted using a sliding scale method with target spectrum width of 0.1 min with a merge tolerance of 100 ppm and maximum retention time gap of 0.5 min.

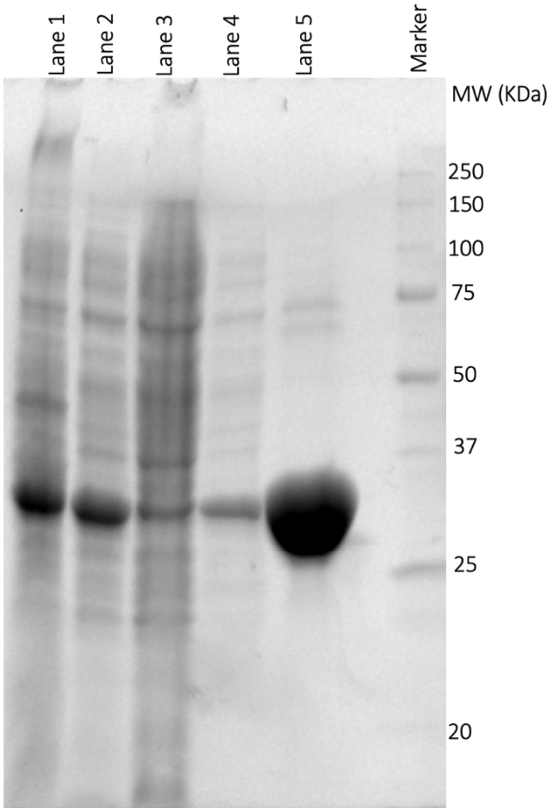

**Figure S4: SDS-PAGE Gel of DhaA31 Purification Fractions**

SDS-PAGE gel depicting the different fractions collected during the purification of DhaA31. Lane 1: Crude cell lysate before induction. Lane 2: Soluble fraction post-sonication. Lane 3: Flow-through from Ni-NTA column. Lane 4: Wash fraction from Ni-NTA column. Lane 5: Eluted fraction containing purified DhaA31. Lane 6: Protein marker (with molecular weights indicated).

A

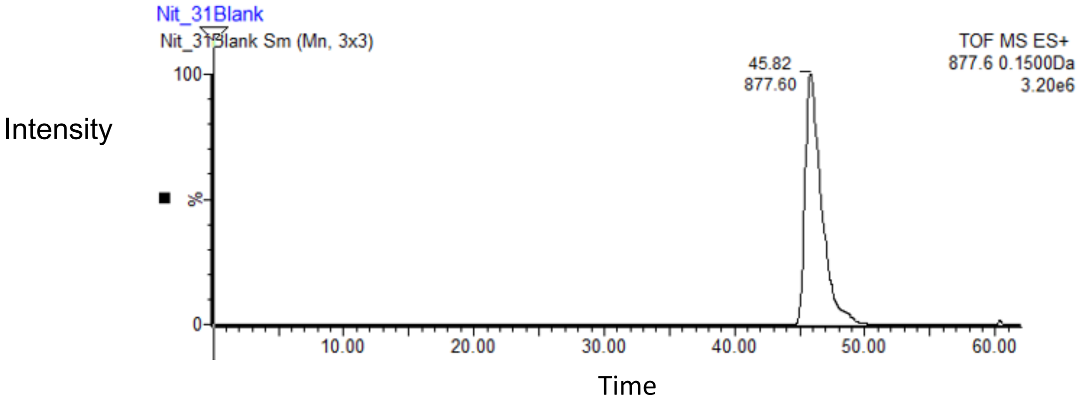

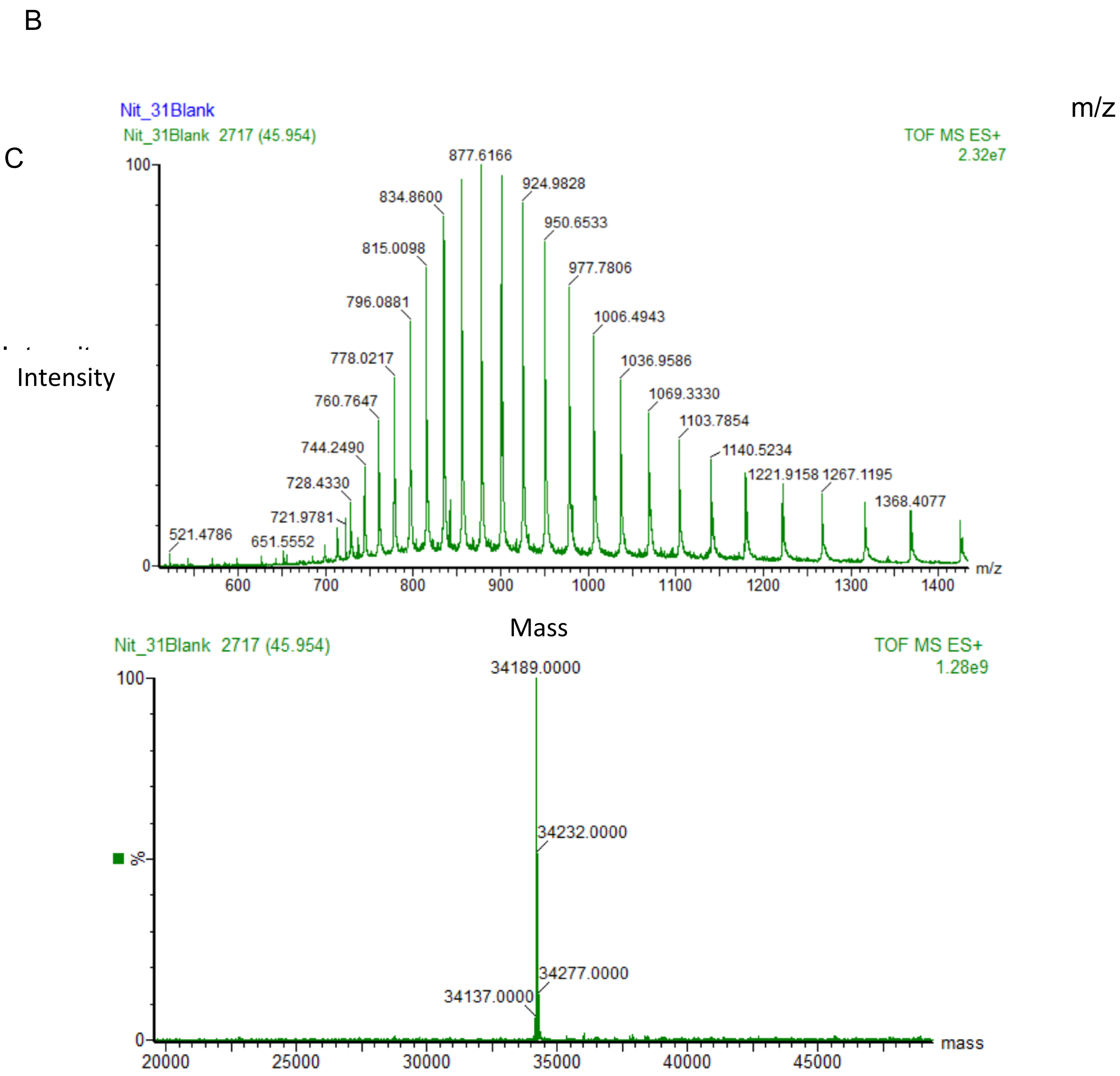


**Figure S5.** LC-MS Analysis of DhaA31 Enzyme. (A)The Intensity vs Time plot displays the extracted ion chromatogram (XIC) for the DhaA31 Enzyme, showing the retention time at approximately 45.82 minutes with a prominent peak at m/z 877.60. The analysis was performed in positive ion mode (ES+). (B) A significant peak at m/z 877.6166 (1500 Da) with an intensity of 2.32e7 is observed, confirming the presence of the DhaA31 Enzyme at this retention time. (C)The deconvoluted mass spectrum of the full protein peak is shown. Calculated Mass 34324Da, however the observed mass 34189.

#### Protein Sequences and Calculated Masses

Table S3

| Protein Sequence [b] | Calculated Mass |
|---|---|
| **MSEIGTGFPFDPHYVEVLGERMHYVDVGPRDG**<br><br>**TPVLFLHGNPTSSYLWRNIIPHVAPSHRCIAPDL** | Expected Mass=34324 Da,<br><br>Observed Mass=34189 Da |

| | |
|---|---|
| **IGMGKSDKPDLDYFFDDHVRYLDAFIEALGLEE**<br><br>**VVLVIHDWGSALGFHWAKRNPERVKGIACMEFI**<br><br>**RPFPTWDEWPEFARETFQAFRTADVGRELIIDQ**<br><br>**NAFIEGALPKYVVRPLTEVEMDHYREPFLKPVD**<br><br>**REPLWRFPNELPIAGEPANIVALVEAYMNWLHQ**<br><br>**SPVPKLLFWGTPGFIIPPAEAARLAESLPNCKTV**<br><br>**DIGPGLHFLQEDNPDLIGSEIARWLPALHHHHHH** | [Expected Mass-Methionine(M)= 34193 Da][a] |

[a]The difference in calculated and observed mass is suspected to be an *N*-terminal processed variant of the desired protein, resulting in removal of the *N*-terminal methionine. [b]Methionine residue highlighted in red.

### Dimer Investigation - Simulation and Calculations

Dehalogenase XRD experiments produced a PDF with a secondary peak at 60 Å. Noting that other features of the PDF match well with simulated calculations, the authors theorized the possibility that this difference in the distributions represented heterogeneity among proteins in the experimental sample, possibly due to dimerization.

A reference AA simulation was performed to investigate this possibility. The initial structure for simulation was derived from PDB 6TY7, a DhaA115 domain-swapped dimer with molecular weight 69.75 kDa, roughly twice the size of the monomer in PDB 3RK4. This structure was simulated for 5ns using the CHARMM36 force field and simulation parameters equivalent to those in the AA simulations of the main manuscript. The simulated structure showed a stable close-packed pairing for the entire trajectory, with the dimer interface remaining nearly the full diameter of either half of the assembly. The final frame of the trajectory is shown in the figure below. Rg = 27 Å.

The dimer PDF was calculated and compared with the main manuscript structures as shown as the purple curve in the figure below.

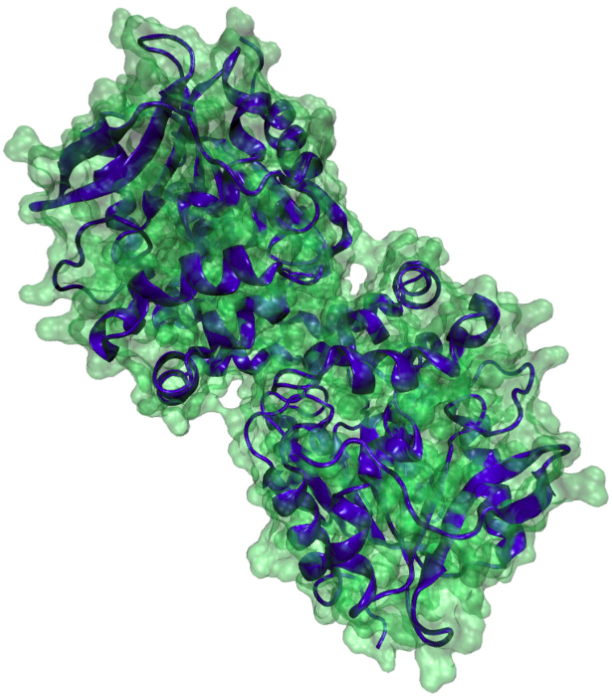

Figure S6. Domain-swapped dimer structure at the end of 5ns simulation. Dimer interface is full diameter of either monomer, creating a nearly cylindrical assembly which persisted over the entire simulated trajectory.

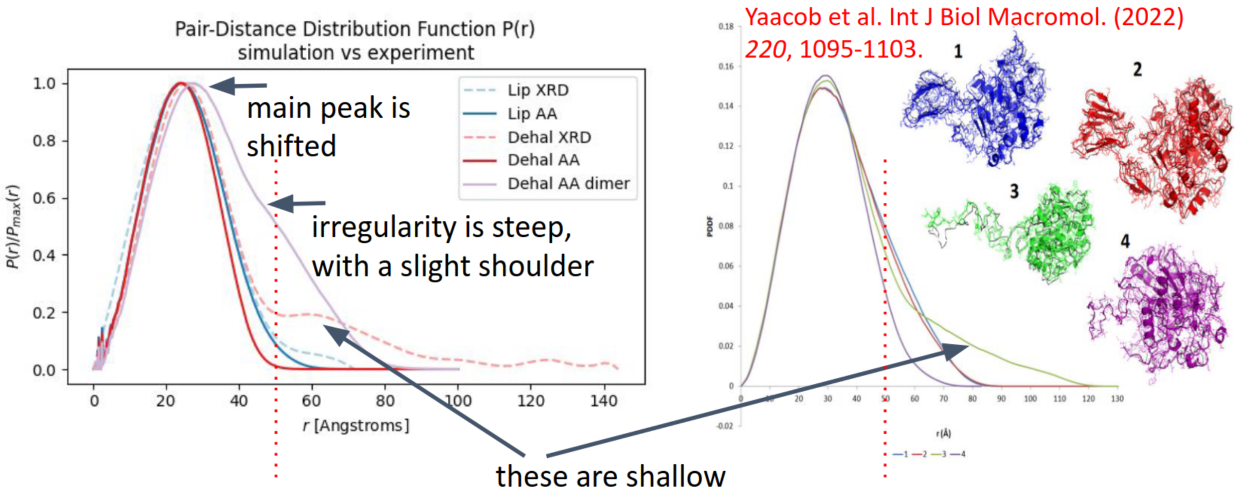


Figure S7. Dimer PDF with comparative graphics copied from Yaacob 2022. On the left, Red Dotted Line is Dhaa PDF via XRD. Long range (50+ Ang) PDF looks like ‘strung out’ green model 3 on the right, a globular protein with a tailing disordered region. Purple line on the left is a simulated dimer.

Further calculations were done using the expressions derived in reference [Oster 1952]. The averaged scattered intensity for a model dimer, two spheres in contact, is

$$I_{dim}(q) = F^2 \frac{1}{4}\left[2 + 2\frac{\sin 2qR}{2qR}\right]$$

Where *F* for a sphere is

$$F = 3\left[\frac{\sin qR - qR\cos qR}{qR^3}\right]$$

In these expressions, *q* is the scattering vector (4p sin(q)/l) and *R* is the radius of the sphere. The intensity for a mixture of monomers and dimer is given by

$$I_{mix}(q) = x I_{mon}(q) + (1-x) I_{dim}(q)$$

Where $I_{mon}(q) = F^2$ and *x* is the monomer fraction

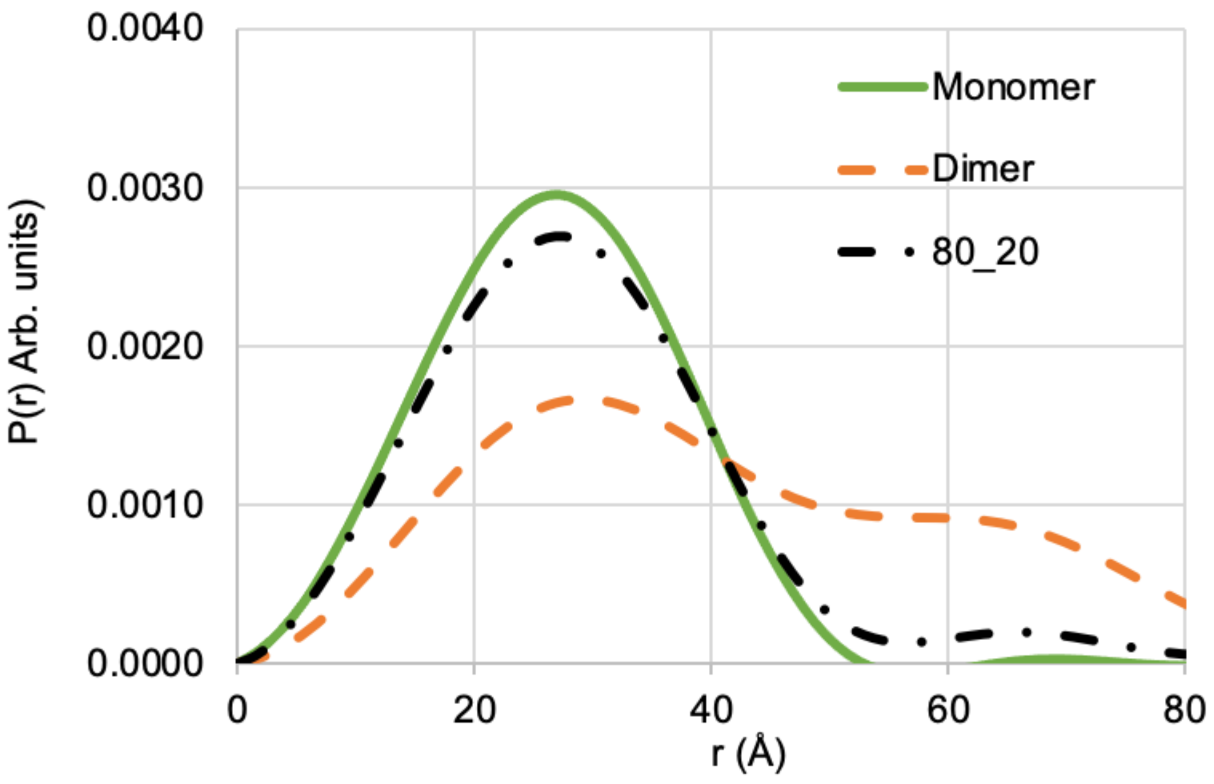


Figure S8. Comparison of the PDFs of a monomer, a dimer, and a mixture with 20% dimer

#### References


Oster, G.; Riley, D. P. Scattering from Isotropic Colloidal and Macro-Molecular Systems. Acta Crystallographica 1952, 5 (1), 1–6. https://doi.org/10.1107/S0365110X52000010.

Yaacob, N.; Kamonsutthipaijit, N.; Soontaranon, S.; Leow, T. C.; Rahman, R. N. Z. R. A.; Ali, M. S. M. Structural Interpretations of a Flexible Cold-Active AMS8 Lipase by Combining Small-Angle X-Ray Scattering and Molecular Dynamics Simulation (SAXS-MD). *International Journal of Biological Macromolecules* **2022**, *220*, 1095–1103. https://doi.org/10.1016/j.ijbiomac.2022.08.145.
